# The oldest fossil record of a living true eel lineage (Protanguillidae, Anguilliformes) from Mexico reveals the gradual acquisition of modern eel traits

**DOI:** 10.1101/2025.09.03.674097

**Authors:** Sebastián Huacuja-Barraza, Kleyton M. Cantalice

## Abstract

We present an integrative taxonomic study describing the first fossil representative of the family Protanguillidae from the Danian deposits near Palenque, Chiapas, Mexico. Osteological analysis and parsimony-based phylogenetic reconstruction using morphological characters support the designation of a new genus and species as a protanguillid by the presence of several features, including the autogenous premaxillae, the presence of metapterygoid and symplectic bones, the participation of pterosphenoid on the posterior margin of orbital margin, the reduced number of vertebrae, and the configuration of hypural complex. The new taxon differs from *Protanguilla palau* by the shape of last branchiostegal ray, the disposition of anterodorsal branch of subopercle, and the insertion of unpaired fins. To investigate the evolutionary history of true eels, we conducted a total-evidence tip-dating analysis combining 14 mitochondrial genes with a morphological matrix, incorporating both Cretaceous and Cenozoic fossil taxa. The results support a gradual acquisition of the diagnostic characters of Anguilliformes throughout the Cretaceous. A Bayesian topological test further supports the species here described as a surviving lineage of Cretaceous eels with †*Libanechelyidae bultyncki*, closely related to the modern eels. This discovery expands the paleobiogeographic range of the Protoanguillidae and provides new insight into the origin and diversification of true eels.

## Introduction

True eels (order Anguilliformes) are a diverse clade of elongated and cylindrical fishes, comprising approximately 1,080 species distributed across 160 genera and 20 families [1,2]. This group is globally distributed, inhabiting a wide range of ecosystems, from coral reefs (e.g., families Muraenidae, Chlopsidae, and Ophichthidae), and freshwater environments (family Anguillidae), to burrowing habits (families Heterenchelyidae and Moringuidae), and abyssal zone (families Saccopharyngidae, Eurypharyngidae, Cyematidae, Monognathidae, and Neocyematidae) [3,4].

Although the anguilliform body plan has evolved convergently in multiple fish lineages, true eels are distinguished by several unique features, including the fusion of the rostral bones into the premaxillo-ethmovomer complex; the bony fusion of the symplectic and metapterygoid bones with the quadrate within the suspensorium; posterior enlargement of the opercular bones; the complete absence of a pelvic girdle; and the continuous integration of the dorsal, anal, and caudal fins [5].

Among extant taxa, *Protanguilla palau* Johnson, Ida & Sakaue, 2012 [6] is unique in retaining several plesiomorphic features otherwise restricted to Cretaceous fossil eels. These features include a free premaxilla; autogenous metapterygoid and symplectic bones; a high concentration of branchiostegal rays the anterior ceratohyal bone in the hyoid arch; a reduced number of vertebrae; unfused hypural plates in the caudal-fin skeleton; and the presence of small imbricated scales [6,7]. Cretaceous eels further retain ancestral traits absent in all modern anguilliforms, such as separate endopterygoid and ectopterygoid bones in the palatoquadrate complex; reduced but present posttemporal bones connecting the pectoral girdle to the skull; and the presence of pelvic fins [8–13].

Some Cretaceous taxa, such as †*Abisaadia* Belouze, 2002 [9], and †*Anguillavus* Hay, 1903 [11], exhibit a suite of fully plesiomorphic characters, while others (e.g., †*Libanechelys bultyncki* Taverne, 2004 [14]) show a mosaic of ancestral and derived traits. In contrast, certain Cretaceous forms like †*Enchelion montimum* Hay, 1903 [11] and †*Nardoechelys robinsi* Taverne, 2002 [15] exhibit derived features typical of modern deep-sea anguilliforms yet retain key ancestral features that support affinities with early-diverging lineages [7,15,16].

The oldest known Cenozoic eels include the enigmatic †*Mylomyrus frangens* Woodward, 1910 [17] and the Monte Bolca eels [18–20], which exhibits the derived traits of modern eels, such as the fusion of the premaxillae with the ethmovomer complex, as well as the formation of a palatopterygoid arch by the palatine, ectopterygoid, and endopterygoid bones, along with components of the suspensorium. Despite these distinct morphological features, the relationships of these taxa to other eels remain unclear. Additionally, there has been described some fossils belonging to extant genera or families [21–25] during Cenozoic, with some otolith records dated back to the Cretaceous [26–30].

While the monophyly of Anguilliformes is well supported by both morphologic and molecular phylogenies [5,31–36], significant discordances persist between these lines of evidence. The case of *P. palau* exemplifies this issue: due its primitive osteology it has been interpreted as a relict lineage of Cretaceous eels [6,37,38], yet molecular analysis consistently places it within Synaphobrachoidei, a clade of bathydemersal modern eels [25,39,40]. Notably, the only phylogenetic analysis to incorporate Cretaceous anguilliforms [9] predates the discovery of †*L. bultyncki* and *P. palau*, limiting its relevance to the current hypothesis.

Here, we describe the first protoanguilid fossil from the Paleocene of Mexico and incorporate both fossil and extant taxa into a comprehensive phylogenetic framework. We employ a total evidence approach and use Bayesian topological tests to evaluate whether protanguillids represent early-diverging members of modern eels or a surviving lineage of stem anguilliforms from the Cretaceous.

### Geological settings

The material was recovered from the Belisario Domínguez quarry, situated approximately 9.5 km south of the city of Palenque, near the village of Belisario Domínguez, Chiapas, Mexico. This locality corresponds to the Tenajapa-Lacandón geological unit [41–43], which has been dated to the early Paleocene (Danian, 63 ± 1.5 Ma) based on strontium isotope analyses conducted on pycnodontiform teeth [44]. The lithology of the outcrops consists primarily of yellow marls and limestone deposited in a shallow marine environment [41–43]. Notably, these sedimentary rocks have a long story of human use, having been employed in the construction of Mayan ceremonial sites and still being utilized by local artisans in contemporary Palenque [41–43].

The Paleocene ichthyofauna of the Belisario Dominguez quarry comprises extinct and extant lineages. It includes extinct representatives of the order Pycnodontiformes and Osteoglossiformes [42], as well as taxa belonging to extant clades, such as Clupeiformes [43] and Gonorynchiformes [45]. Members of acanthomorph groups are also present, including Serranidae [46], Pomacentridae [47], undetermined percomorphs [48], and Syngnathiformes [49]. In addition to these fishes, the assemblage contains fragmentary remains of other vertebrates and terrestrial plant material [42,43]. Eel-like fossils were previously recognized at the site and tentatively assigned to Anguilliformes based on the presence of key synapomorphies, however, no formal taxonomic identification was proposed at time [42].

## Material and methods

A total of 22 specimens in several states of preservation were examined for this study. All specimens are housed in the *Colección Nacional de Paleontología* of the *Instituto de Geología* at the *Universidad Nacional Autónoma de México*, Mexico City, under accession numbers IGM1300 to IGM1316. Mechanical preparation was carried out on select specimens using needles and micro-excavators to carefully remove matrix material obscuring anatomical features. One specimen (IGM1305) was transferred to polyester resin and the matrix was dissolved using acid following the method of Toomb and Rixon [50] to facilitate detailed morphological analysis

Specimens were photographed using a Nikon D5600 DSLR with AF-S Micro NIKKOR 60 mm macro lens under both incandescent and ultraviolet light; Microphotographs were obtained with a ZEISS Axio Zoom v16 stereomicroscope and processed using Zen 3.5 Blue Edition software for image stacking and enhancement.

Anatomical observations were conducted under a stereomicroscope, and linear measurements were taken using a digital caliper, following the osteological terminology proposed by McCosker [51]. Owing to the poor preservation of the opercular bones (often fragmented and disarticulated), cephalic length was replaced with cranial length, defined as the distance from the anterior tip of the rostrum to the posterior margin of supraoccipital. Comparative morphological data were obtained from previously published sources [7,52–60].

### Nomenclatural acts

The electronic edition of this article conforms to the requirements of the amended International Code of Zoological Nomenclature, and hence the new names contained herein are available under that Code from the electronic edition of this article. This published work and the nomenclatural acts it contains have been registered in ZooBank, the online registration system for the ICZN. The ZooBank LSIDs (Life Science Identifiers) can be resolved and the associated information viewed through any standard web browser by appending the LSID to the prefix “http://zoobank.org/”. The LSID for this publication is: urn:lsid:zoobank.org:pub:114D2528-7A15-4318-9932-0B584463D934. The electronic edition of this work was published in a journal with an ISSN and has been archived and is available from the following digital repositories: PubMed Central, LOCKSS.

### Abbreviations

**IGM**, *Instituto Geológico de México* [Mexican Institute of Geology], is the catalog registration abbreviation for specimens deposited in the Colección Nacional de Paleontología [National Collection of Paleontology of Mexico]. IGM loc. is the catalog abbreviation for fossil localities on the National Collection of Paleontology of Mexico.

### Phylogenetic analysis

#### Morphological analysis

Morphology-based phylogenetic analysis was performed using the matrix originally proposed by Belouze [7], which comprises 123 characters and 61 taxa, including 40 extant and 21 fossils species. The matrix was modified using Mesquite v.3.70 [61]. The genus *Osteoglossum* was excluded due to its unstable phylogenetic placement in the original analysis [7], and *Tarpon* was removed due to its synonym with *Megalops* [62]. Newly incorporated taxa include *Protanguilla palau*, †*Nardoechelys robinsi* and †*Libanechelys bultyncki,* and the species described herein. Characters for these taxa were coded based on original descriptions and subsequent anatomical assessments [6,14–16,37,38]. The resulting dataset comprises 62 taxa and retains the original set of 123 characters.

Parsimony analyses were performed using TNT v. 1.6 [63]. A heuristic search was conducted using *Hiodon* designed as the outgroup. The search began with a Wagner tree using a random seed of 1 and employed the Tree Bisection and Reconnection (TBR) algorithm. The analysis was run for 1,000 replicates, saving up to 10 trees per interaction. Two datasets were analyzed: (1) the complete matrix including all 62 taxa, and (2) a reduced matrix excluding all Cenozoic fossil taxa, except for the new species described herein. The exclusion of certain Cenozoic taxa was based on their unstable placement in preliminary analyses, likely resulting from a combination of fossil incompleteness, high levels of homoplasy within the group, and uncertain taxonomic assignment. We also carried a Bootstrap analysis with 1000 generations in both matrixes.

In addition, New Technology searches in TNT were used to assess the potential for recovering additional topologies. However, no further most parsimonious trees (MPTs) were obtained through these search strategies.

#### Total evidence analysis

To compare the results of the morphology-based parsimony analysis with a parametric approach integrating molecular data, we retrieved the mitogenomic sequences from GenBank for one representative species of each genus included in the original morphological matrix (Table 1). Due to the unique mitochondrial rearrangements observed in “Saccopharyngoidei” [64,65] we extracted 14 individual protein-coding and ribosomal genes based on GenBank annotations using Geneious software and aligned the sequences with MAFFT v. 7.526 [66], employing default settings.

**Table 1.** Sequences retrieved from GenBank for each taxon.

| **Familiy** | **Species** | **Data Accession** |
| --- | --- | --- |
| Albulidae | *Albula vulpes* | OR546140.1 |
| Anguillidae | *Anguilla anguilla* | KJ564270.1 |
| Chlopsidae | *Chilorhinus platyrhynchus* | OP035172.2 |
| Colocongridae | *Coloconger cadenati* | AP010863.1 |
| Congridae | *Ariosoma meeki* | OK585090.1 |
| Congridae | *Heteroconger hassi* | NC_013629.1 |
| Cyematidae | *Cyema atrum* | AP010870.1 |
| Derichthyidae | *Derichthys serpentinus* | AP010851.1 |
| Elopidae | *Elops saurus* | NC_005803.1 |
| Eurypharyngidae | *Eurypharynx pelecanoides* | AB046473.2 |
| Halosauridae | *Halosaurus ovenii* | MW077726.1 |
| Heterenchelyidae | *Pythonichthys microphtalmus* | NC_013601.1 |
| Hiodontidae | *Hiodon alosoides* | AP004356.2 |
| Megalopidae | *Megalops atlanticus* | AP004808.1 |
| Moringuidae | *Moringua microchir* | NC_013602.1 |
| Muraenesocidae | *Cynoponticus ferox* | AP010853.1 |
| Myrocongridae | *Myroconger compressus* | NC_013631.1 |
| Nemichthyidae | *Facciolella oxyrhyncha* | AP010866.1 |
| Nemichthyidae | *Hoplunnis punctata* | AP010865.1 |
| Nemichthyidae | *Nemichthys scolopaceus* | AP010854.1 |
| Nemichthyidae | *Nettastoma parviceps* | AP010864.1 |
| Notacanthidae | *Notacanthus bonaparte* | MT027058.1 |
| Notacanthidae | *Polyacanthonotus caballeroi* | KT592358.1 |
| Ophichthidae | *Myrophis microchir* | NC_083176.1 |
| Ophichthidae | *Ophichthus erabo* | OP154196.1 |
| Protanguillidae | *Protanguilla palau* | AP011809.1 |
| Saccopharyngidae | *Saccopharynx lavenbergi* | AP018343.1 |
| Serrivomeridae | *Serrivomer sector* | AP007250.1 |
| Synaphobranchidae | *Ilyophis brunneus* | NC_013634.1 |
| Synaphobranchidae | *Simenchelys parasitica* | AP010849.1 |
| Synaphobranchidae | *Synaphobranchus oregoni* | OP056972.1 |

Mitogenomic sequences are retrieved from GenBank with taxonomic specifications for family and species.

Following manual revision and correction, sequence alignments were analyzed in Partition Finder v. 2.1.1 [67] to identify the optimal partitioning scheme and molecular evolutionary models (Table 2) under the Akaike Information Criterion (AIC). Morphological data were analyzed separately using the MKv model [68]. Subsequently, the molecular and morphological datasets were concatenated into a single matrix, excluding Cenozoic fossil taxa (resulting in a total of 42 taxa).

**Table 2.**
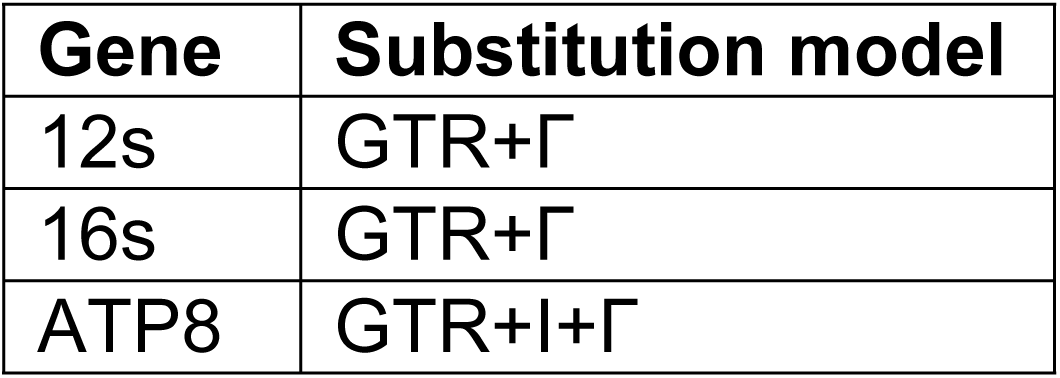
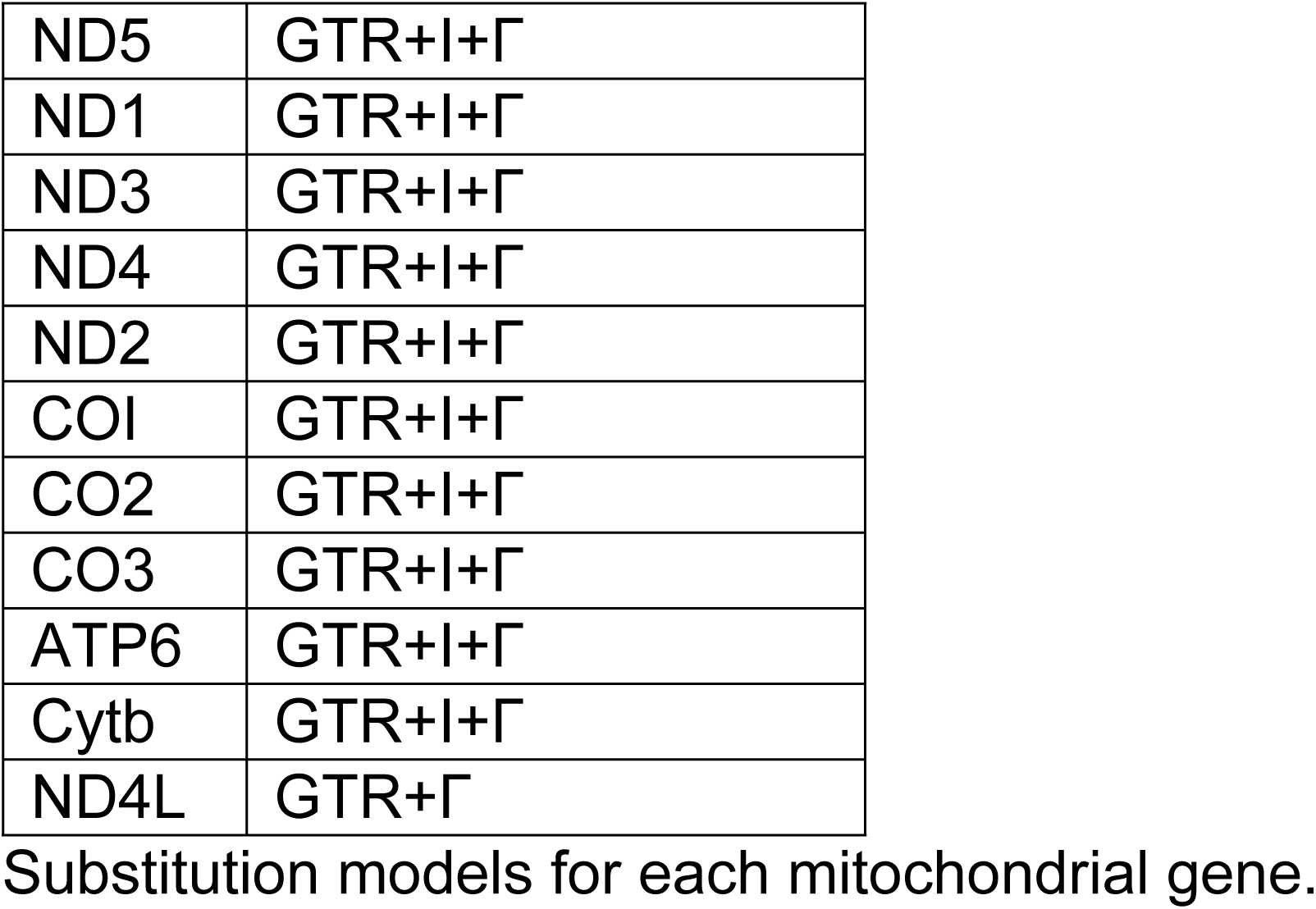
Substitution models for each gene.

A tip-dating (TD) analysis was performed in MrBayes v. 3.2 [69], incorporating nine fossil taxa in addition to a root calibration derived from the oldest confidently identified elopomorph fossil [70,71] under the fossilized birth-death (FBD) process. A preliminary analysis of the sequence matrix was conducted to estimate molecular clock parameters. Fossil ages priors were modeled using a uniform distribution (Table 3), informed by stratigraphy information obtained from the Paleontology Database (https://paleobiodb.org/#/), as well as original descriptions [14] and taxonomic revisions [7–9,42]. Furthermore, topological constraints were applied to Cretaceous fossil taxa and *P. palau* based on the results of the parsimony analysis.

**Table 3.** Fossil ages used in the tip-dating analysis. Fossil calibrations used in the tip-dating analysis. Minimum and maximum bounds (in millions of years) were implemented as uniform priors.

| Taxa | Min. age | Max. age |
| --- | --- | --- |
| root | 140 | 150 |
| *Enchelurus* | 72.2 | 100.5 |
| *Abisaadia hakelensis* | 93.9 | 100.5 |
| *Anguillavus* | 93.9 | 100.5 |
| *Luenchelys minor* | 93.9 | 100.5 |
| *Hayenchelys germanus* | 93.9 | 100.5 |
| *Urenchelys avus* | 83.6 | 100.5 |
| *Libanechelys bultyncki* | 93.9 | 100.5 |
| †*Mitsijt tsucul* gen. et sp. nov. | 61.5 | 64.5 |

Eight independent Markov chain Monte Carlo (MCMC) chains were run for 100 million generations each, with sampling conducted every 1,000 generations. A 25% burn-in was applied prior to tree summarization. Convergence diagnostics were evaluated in Tracer v. 1.7 [72], with Effective Sample Size (ESS) values >200 considered indicative of adequate sampling and convergence of the posterior distributions.

To evaluate whether Protanguillidae or †Libanechelyidae constitutes the sister group of all extant eels (”modern” Anguilliformes), we conducted hypothesis testing using both the morphological and total-evidence datasets (Fig. 1). Marginal likelihood was estimated via stepping-stone sampling (SS test) [73] implemented in MrBayes, employing the same model parameters and topological constraints in the tip-dating analyses.

**Figure 1.**
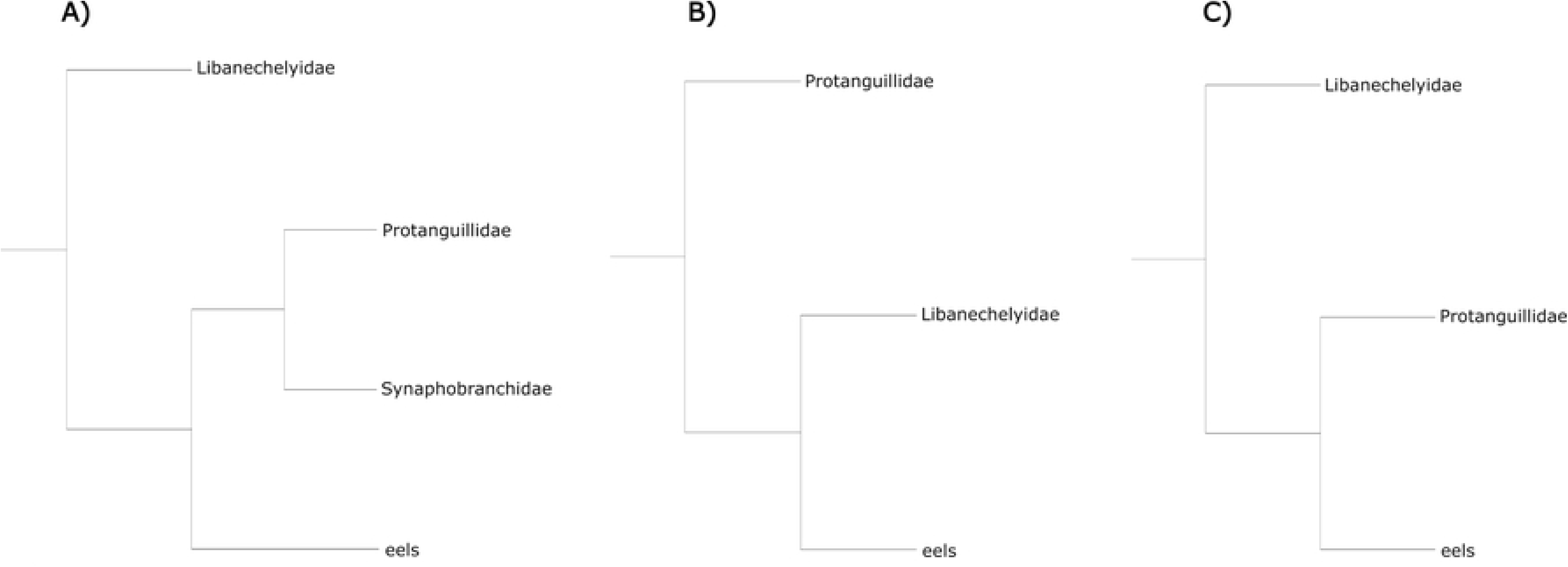
Topologies tested for the sister group of the modern eels. We estimated marginal likelihoods using the stepping-stone method for (A) the topology recovered in the total evidence analysis, (B) the Protanguillidae hypothesis obtained from the morphological analysis, and (C) the †Libanechelyidae hypothesis as proposed by Taverne [14].

## Results

### Systematic Paleontology

Division Teleostei Müller, 1845 [74]

Cohort Elopomorpha Greenwood, Rosen, Weitzman & Myers, 1966 [75]

Order Anguilliformes Goodrich, 1909 [76]

Family Protanguillidae Johnson, Ida & Sakaue, 2012 [6]

†*Mitsijt* gen. nov.

### Type species

†*Mitsijt tsucul* gen. et sp. nov.

### Diagnosis

As the species described below.

### Etymology

The generic name *Mitsijt* is derived from the Ch’ol word mits’ijt, meaning “eel.” Although the term traditionally refers to mud eels of the order Synbranchiformes, it is also used locally to describe both living and fossilized fishes exhibiting a cylindrical and elongated body shape.

†*Mitsijt tsucul* sp. nov.

### Holotype

IGM1300, 174 mm of SL. A specimen preserved on its left side, showing an autogenous premaxilla, symplectic and metapterygoid bones, expanded 11^th^ branchiostegal ray broader than subopercle and spinous complex of the anteriormost vertebrae.

### Paratypes

IGM1308, 91.2 mm of SL, preserved on its right side showing the suspensorium bones *in situ*; IGM1311, 123 mm of SL, preserved in dorsal view showing the anterior part of ethmovomer complex and skull roof; IGM1314, incomplete, preserved in ventral view showing the hyoid arch; IGM1318, 134 mm of SL, has a good cranial preservation, the different vertebral regions and the best caudal fin complex.

### Referred specimens

22 specimens, recorded as IGM1300-1321.

### Type locality and horizon

Belisario Domínguez quarry (17°25′60″ N, 91°58′46.80″ W,), Salto del Agua Municipality, state of Chiapas, recorded as IGM loc 3870, corresponding to the Tenejapa-Lacandón geological unit dated in the Paleocene epoch, Danian age (63 Ma, ±1.5 Ma) [42].

### Diagnosis

†*Mitsijt tsucul* gen. et sp. nov. is diagnosed by a distinctive combination of morphological characters not observed in any other known Anguilliformes. These include an autogenous premaxilla; the extension of the pterosphenoid into the posterior orbital region; unfused symplectic and metapterygoid bones relative to the quadrate; and the presence of twelve branchiostegal rays, with the terminal ray being laterally expanded and broader than the subopercle. This species exhibits an average of 90 vertebrae (ranging from 83 to 94), with the anterior six neural arches remaining free from their respective vertebral centra, forming a reinforced and spinous process complex. In the caudal skeleton, the ural vertebrae are fused to the ventral hypural plates I–II, while the dorsal hypural plates III–IV remain separate from the compound complex.

### Etymology

The specific name *tsucul* derives from the Ch’ol word for “old” or “ancient”. The generic plus specific name refers to “ancient eel”.

## Description

### General body features

†*Mitsijt tsucul* is an elongated and cylindrical anguilliform fish ranging from 91.2 227 mm in standard length (SL). The cranial length is approximately 9.7% of the SL. The anterior border of the head is triangular. The pectoral fins are vertically positioned and inserted at 13.2% of the SL. No pelvic girdle is observed. The dorsal fin originates at the first third of the body (mean 36.2% of SL) and the anal fin is inserted slightly posteriorly (mean 45.8% of SL), both fins are completely confluent with the anal fin (Tab. 4).

**Table 4.**
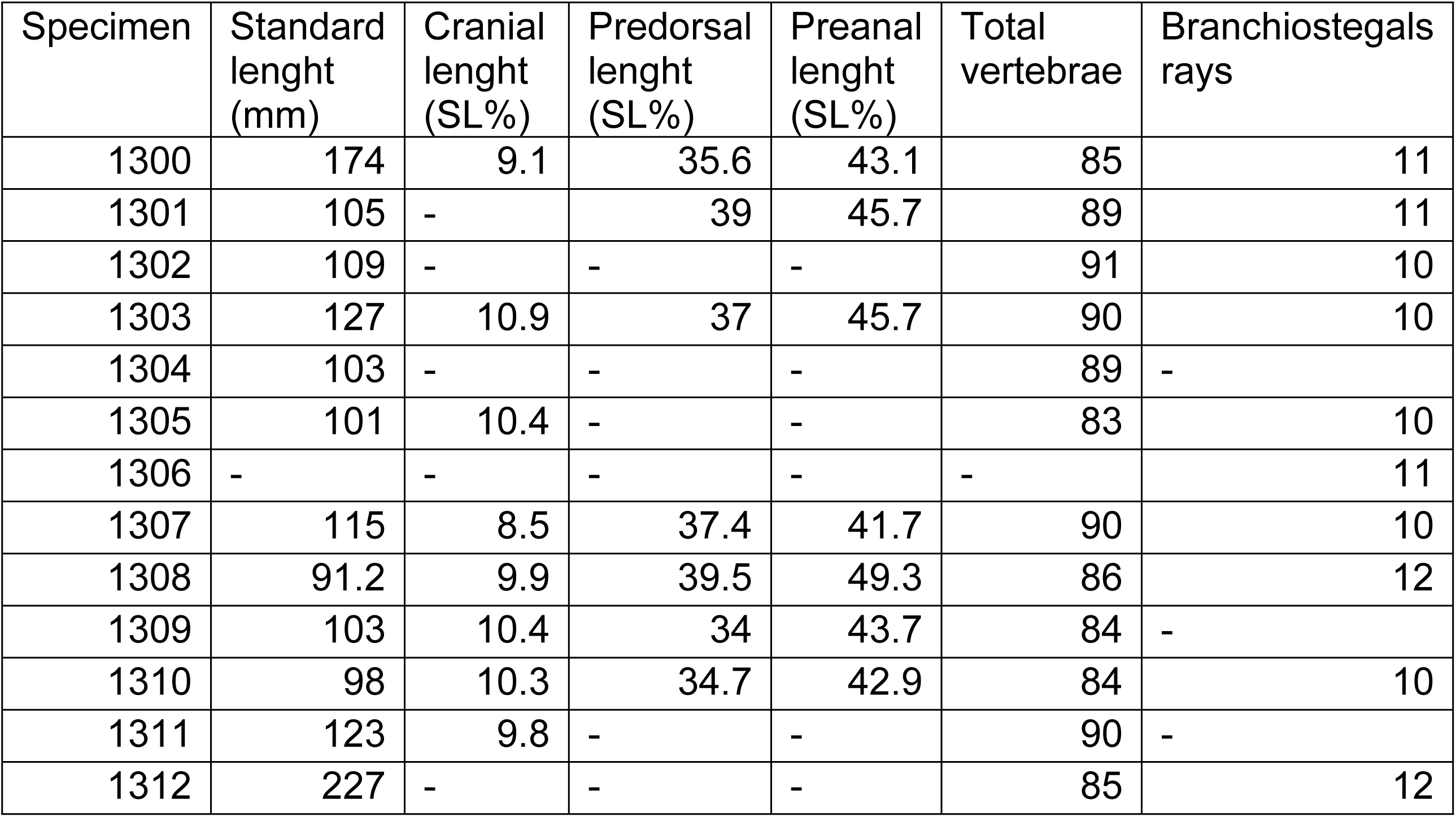
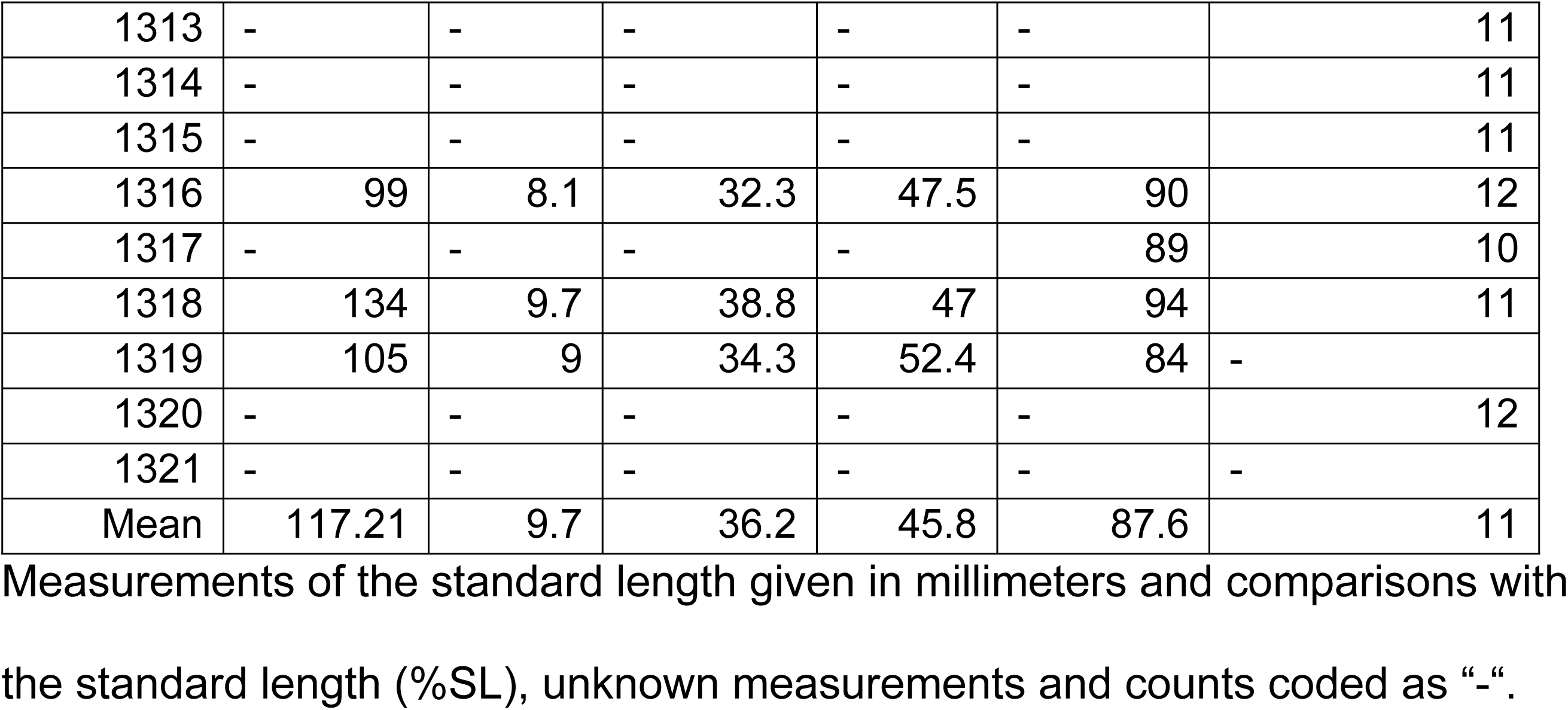
Measurements and meristic counts of the specimens of *M. tsucul*.

### Neurocranium

The anteorbital region is composed of the ethmovomer (**et)**. This bone has a fan-shaped anterior end with numerous sensory pores. It extends posteriorly as a thin bar without any lateral expansion (Fig. 2). The frontals (**f)** are paired and contact the ethmovomer at mid-orbit in a “V” shape. In the postorbital region, these bones expand laterally but are excluded from the cranial margin by the pterotic. In the lateral view, the frontals form at least one-third of the posterior orbital margin.

**Figure 2.**
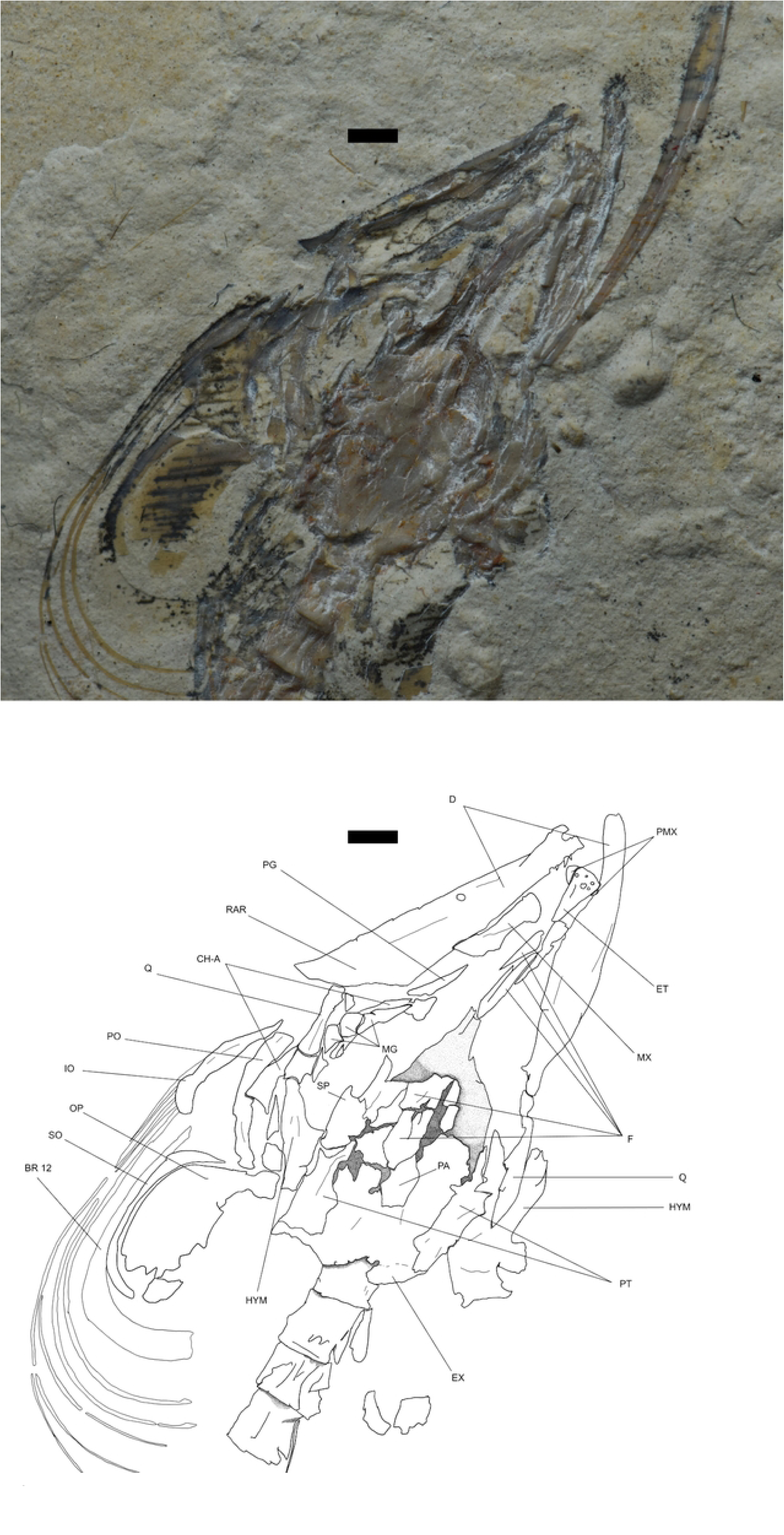
†*Mitsijt tsucul* gen. et sp. nov specimen IGM1311. Dorsal view of IGM1311. Scale bar 0.5 mm.

The parietals (**pa**) are quadrangular, paired bones without any noticeable modifications. The supraoccipital (**soc**) is a triangular bone inserted into the intraparietal suture and extends beyond the level of the epioccipitals (**eo**), which articulate laterally with the pterotic (**pt**) and posteriorly with the semilunar-shaped exoccipitals (**ex**) (Fig. 3).

**Figure 3.**
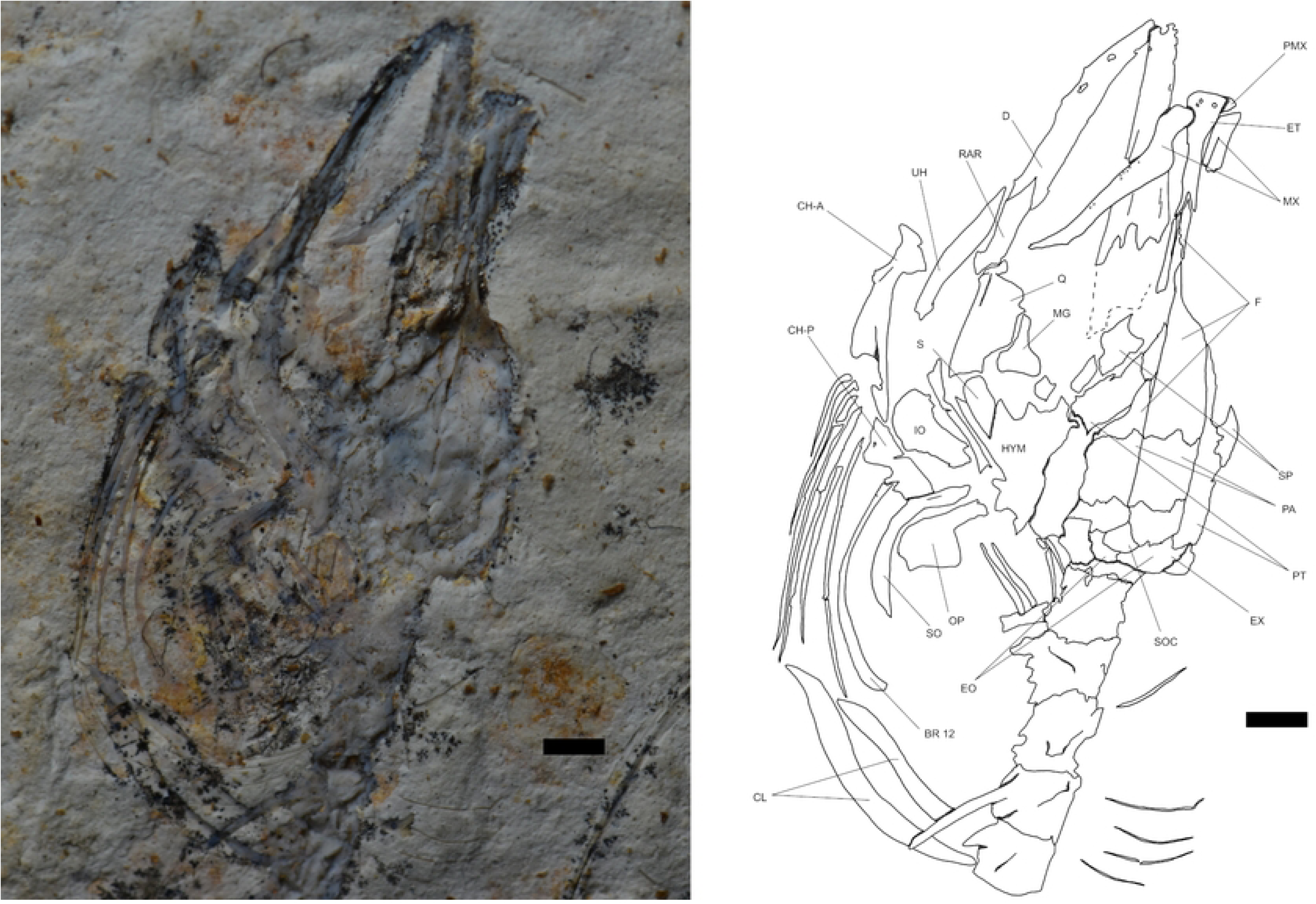
†*Mitsijt tsucul* gen. et sp. nov specimen IGM1316. Dorsal view of IGM1316. Scale bar 0.5 mm.

The pterosphenoid (**pts**) is a well-developed triangular bone that participates in the posterior orbital margin and articulates ventrally with a small triangular basisphenoid (**bs**) (Fig. 4). Ventrally, the ethmovomer articulates with the parasphenoid (**pas**), a slender bone that runs along the neurocranium and connects with the basisphenoid.

**Figure 4.**
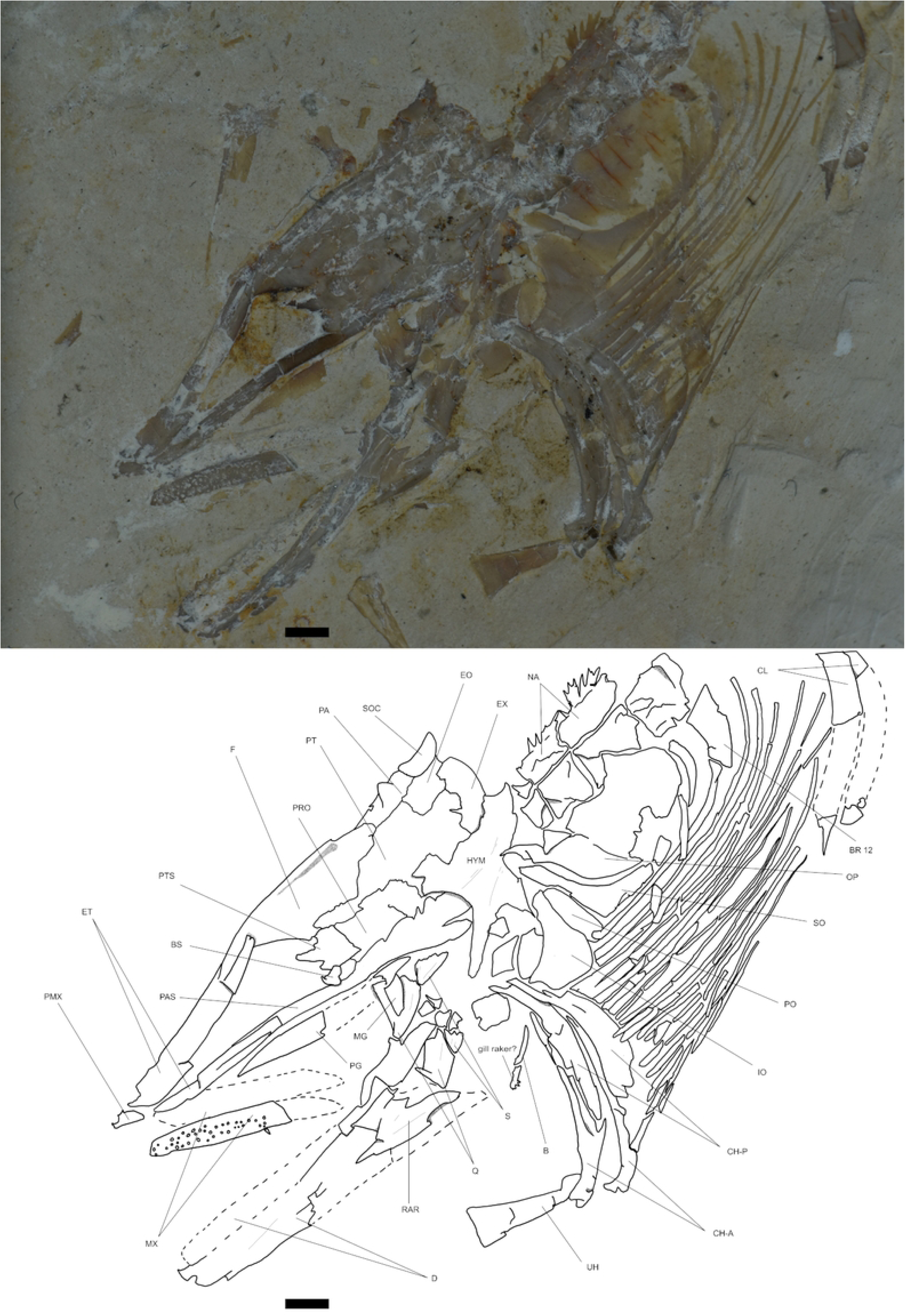
†*Mitsijt tsucul* gen. et sp. nov holotype specimen IGM1300. Lateral view of IGM1300. Scale bar 0.5 mm.

As in other anguilliforms, the pterotic (**pt**) has an anterior process that contacts the pterosphenoid. The posterior body of the pterotic is quadrangular, and the pterotic sensory canal is open and exposed on the medial surface, articulating with the exoccipitals. The sphenoid (**sp**) articulates with the anterior part of the quadrangular body of the pterosphenoid and has a horn-like shape. The prootic (**pro**) articulates anteriorly with the pterosphenoid and ventrally with the parasphenoid. It has a rectangular shape and is the largest lateral component of the neurocranium, but it is excluded from the margin by the basioccipital (**bo**). However, no ventral view of the neurocranium is well preserved, and the basioccipital is only known from lateral view.

### Hyo-palatine bones

As in *P. palau*, the symplectic (**s**) and metapterygoid (**mg**) are independent from the quadrate (**q**). Both are thin, laminar bones that are frequently fragmented or absent. The metapterygoid is a pyramidal bone that articulates with the anterior face of the quadrate, while the symplectic is an elongated, kidney-shaped bone positioned along the posterior face of the quadrate (Fig. 5). The quadrate itself is fan-shaped, with a spherical head that articulates ventrally with the articulo-anguloretroarticular (**rar**).

**Figure 5.**
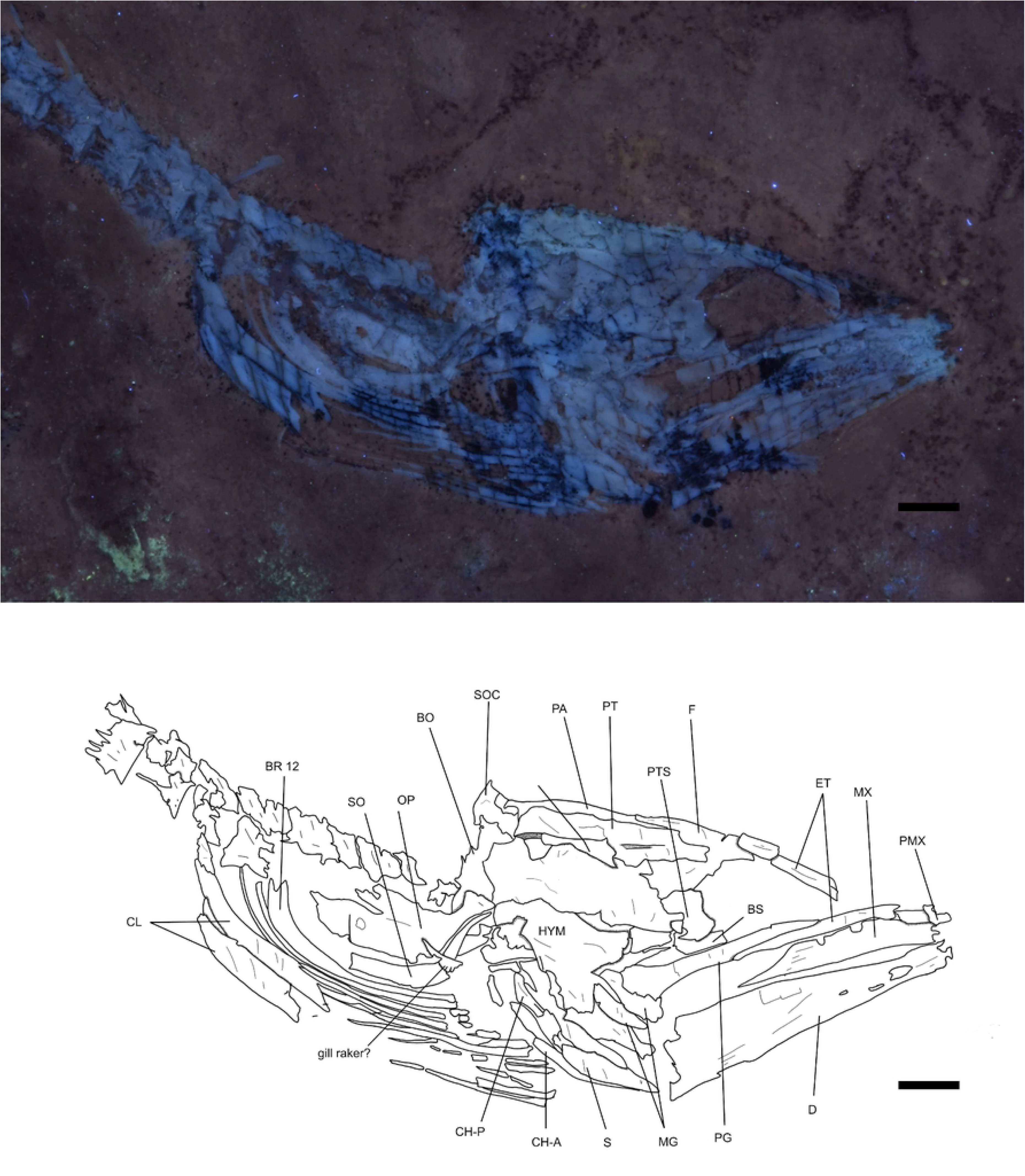
†*Mitsijt tsucul* gen. et sp. nov paratype specimen IGM1308 under UV light. Lateral view of IGM1308. Scale bar 0.5 mm.

The hyomandibular (**hym**) is a robust rhomboid bone with a short posterodorsal projection that articulates with the opercle (**op**). Ventrally, it features two triangular projections that interlock with the quadrate and are supported by a pronounced posterodorsal ridge, like the configuration observed in *P. palau*.

The palatopterygoid arch (**pg**) is slender and delicate, often fragmented or absent in most specimens. Its anterior tip is free and pointed, running along the superior border of the mouth and articulating with the hyomandibular via a broadened posterior end. In ventral view, the arch appears slightly recurved along its medial section.

### Jaws

This eel possesses a free premaxilla (**pmx**) (Fig. 6), a unique characteristic among Anguilliformes observed only in Cretaceous fossils, protanguillids, and *Derichthys* Gill, 1884 [77]. The bone is small and resembles that of *P. palau* in lateral view. In most specimens preserved in dorsal or ventral positions, it is partially covered by the ethmovomer.

**Figure 6.**
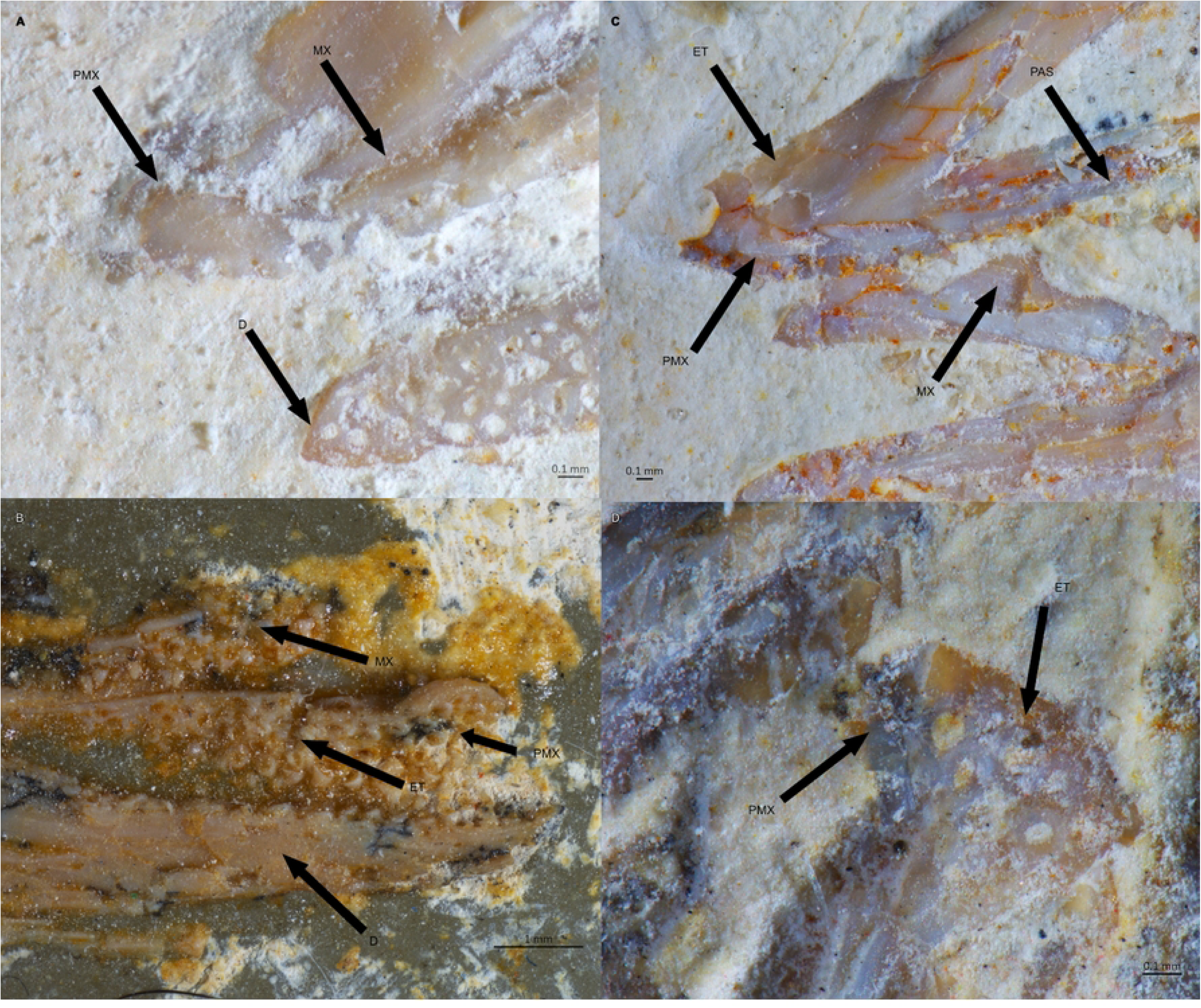
†*Mitsijt tsucul* gen. et sp. nov premaxillae. View of the ethmoidal region of A) holotype IGM1300, B) paratype IGM1318, C) transferred specimen IGM1312 and D) paratype IGM1311. Scale bar 0.1 mm.

The maxilla (**mx**) is slender and displays a distinct, well-developed anterodorsal process. Shortly after, it narrows at a constriction before forming a posterior peduncle that becomes progressively slender, terminating in a pointed margin. The teeth are conical and arranged in biserial rows; in most specimens, they are absent, but the dental sockets are preserved.

The dentary (**d**) is slightly curved anteriorly and features an anterior pore and multiple lateral pores associated with cranial nerves. The coronoid process is poorly developed, while the posterior margin forms an angular shape. Like the maxilla, the dentary’s teeth are conical and arranged in a biserial pattern, although they are only preserved in a few specimens.

As in other anguilliforms, the posterior elements of the lower jaw are fused into a single bone, the articulo-anguloretroarticular. This bone is wedge-shaped, with a pointed anterior end that slides into the dentary.

### Opercular bones

The opercular series is present and posteriorly displaced from the skull, following the typical arrangement observed in anguilliforms. The bones are thin and, in most cases, fragmented or missing. The preopercle (**po**) is narrow and curved, with a posterior expansion forming a boomerang shape. The interopercle (**io**) is as long as the preopercle but thinner and lacks any posterior expansion.

The opercle is oval-shaped, caudally elongated, and features a short, bottleneck-like articular process. It lacks ornamentation or significant modifications. The subopercle (**so**) exhibits a small, dorsally directed anterior process and extends along the ventral edge of the opercle (Fig. 7).

**Figure 7.**
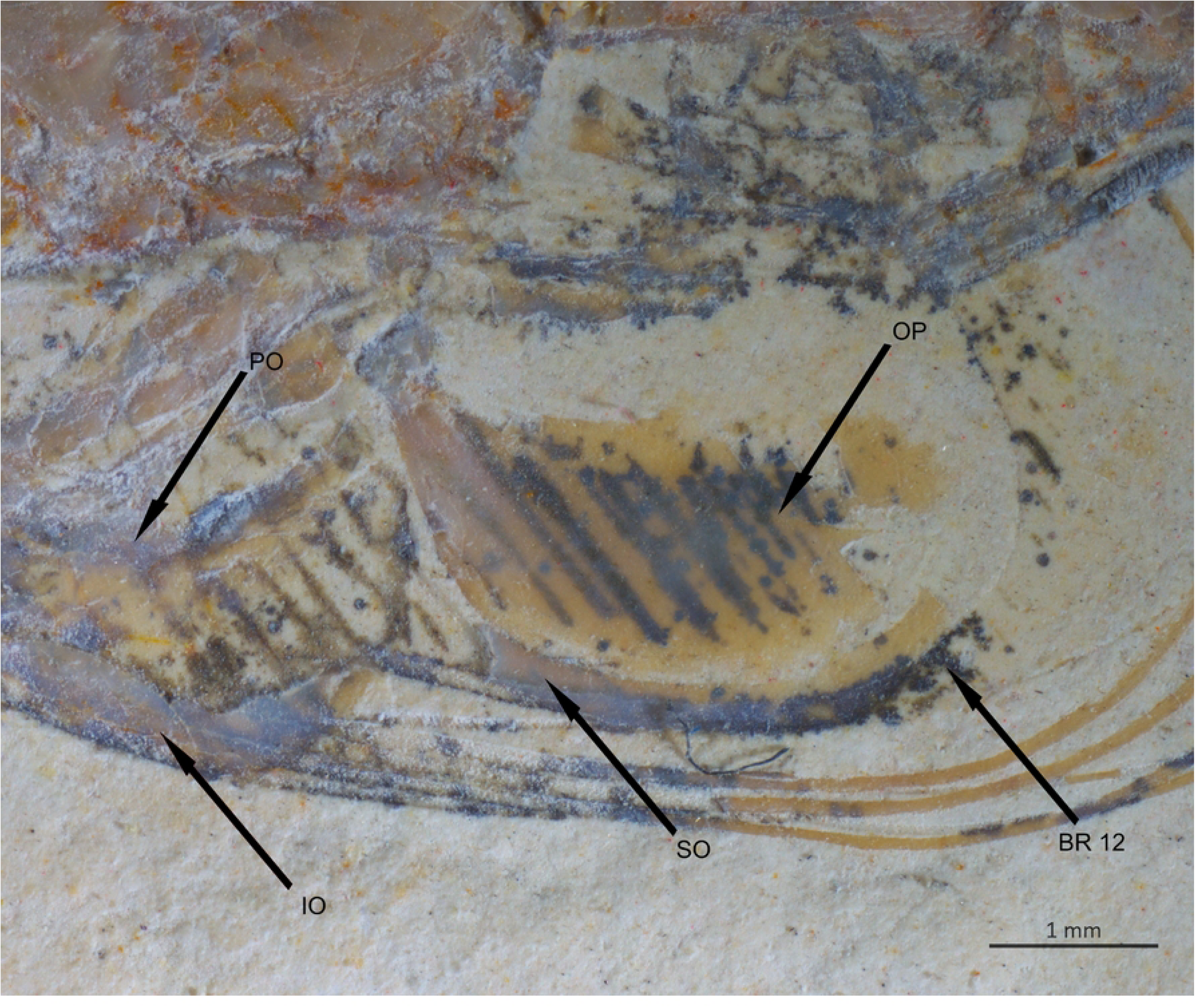
†*Mitsijt tsucul* gen. et sp. nov specimen IGM1311. Dorsal view of IGM1311. Scale bar 0.1 mm.

### Hyoid arch and branchiostegal rays

No urohial is preserved in the specimens, and only a few exhibit a glossohial (**uh**), which appears as a bar that is slightly expanded posteriorly.

The ceratohyals are curved and posteriorly displaced. The anterior ceratohyal (**ch-a**) possesses a ventral hammerhead-shaped region and a long branch that runs along the anterior edge of the posterior ceratohyal (**ch-p**). The posterior ceratohyal is slightly curved and as robust as the anterior one (Fig. 8).

**Figure 8.**
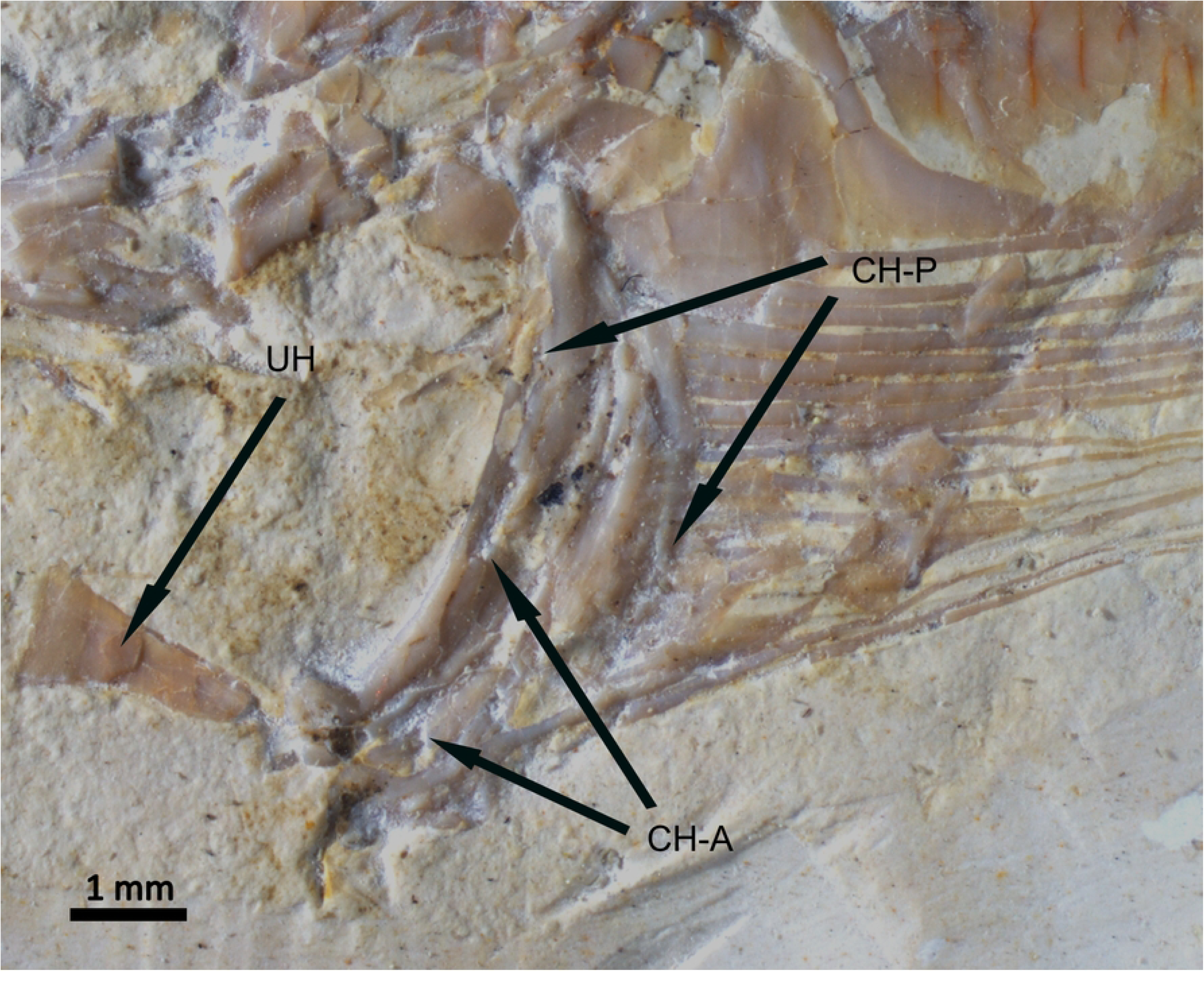
†*Mitsijt tsucul* gen. et sp. nov holotype specimen IGM1300. Lateral view of IGM1300. Scale bar 0.5 mm.

The branchiostegal rays (**br**) are associated with the posterior ceratohyal, with only one or two originating from the anterior ceratohyal. These rays are long, narrow, and curve dorsally towards the opercle. The last branchiostegal ray is significantly expanded from its middle portion; in the largest specimens, it becomes wider than the subopercle.

Some specimens display poorly ossified elements resembling gill arches, but these appear as isolated structures without clear articulation.

### Pectoral girdle and fin

The pectoral fin is well-developed and positioned medially, with the cleithrum (**cl**) being robust and “L”-shaped, present in all specimens. However, the remaining elements of the girdle are poorly preserved, with only a few specimens showing additional structures. The supracleithrum (**scl**) is narrow and curved, while the coracoid (**co**) and scapula (**sc**) are quadrangular and poorly ossified (Fig. 9).

**Figure 9.**
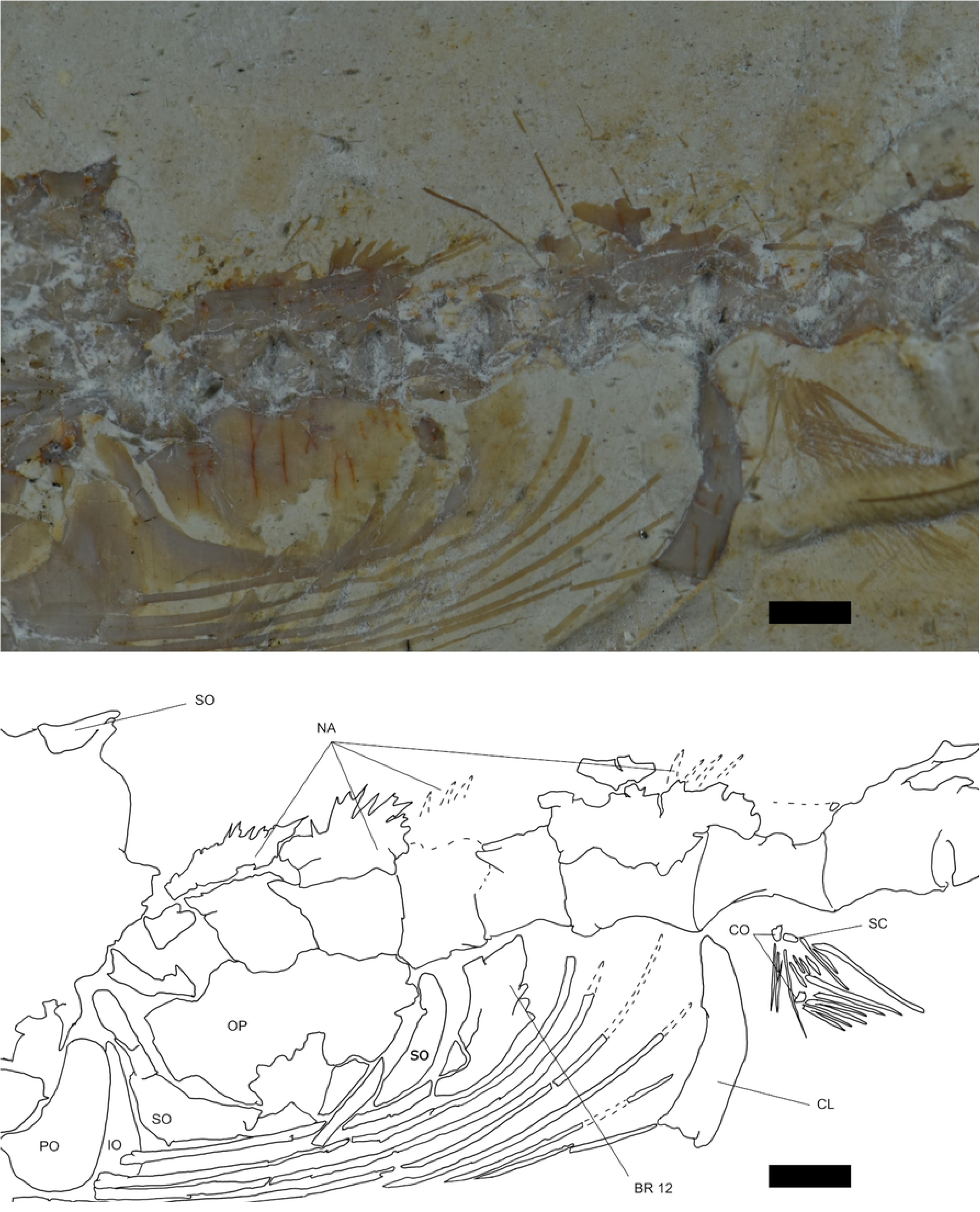
†*Mitsijt tsucul* gen. et sp. nov. holotype specimen IGM1300. Lateral view of IGM1300. Scale bar 0.5 mm.

### Vertebral column, intramuscular bones, dorsal, and anal fin

The six anteriormost vertebrae bear modified neural arches that form a spinous complex. This complex is free from the centrum and fused into a single plate. While the spinous complex itself is not preserved in the specimens, the plate is consistently present (Fig. 9).

The abdominal vertebrae exhibit a parapophysis that is sharply pointed, gradually becoming more robust in subsequent vertebrae until the insertion of the first pterygiophore, and sometimes a little triangular anterior apophysis is also present in the biggest specimens. In the caudal vertebrae (those associated with the unpaired fins), the hemal arches adopt a saddle shape with strong, posteriorly directed hemal spines. In some specimens, a small anterior projection is also observed (Fig. 10).

**Figure 10.**
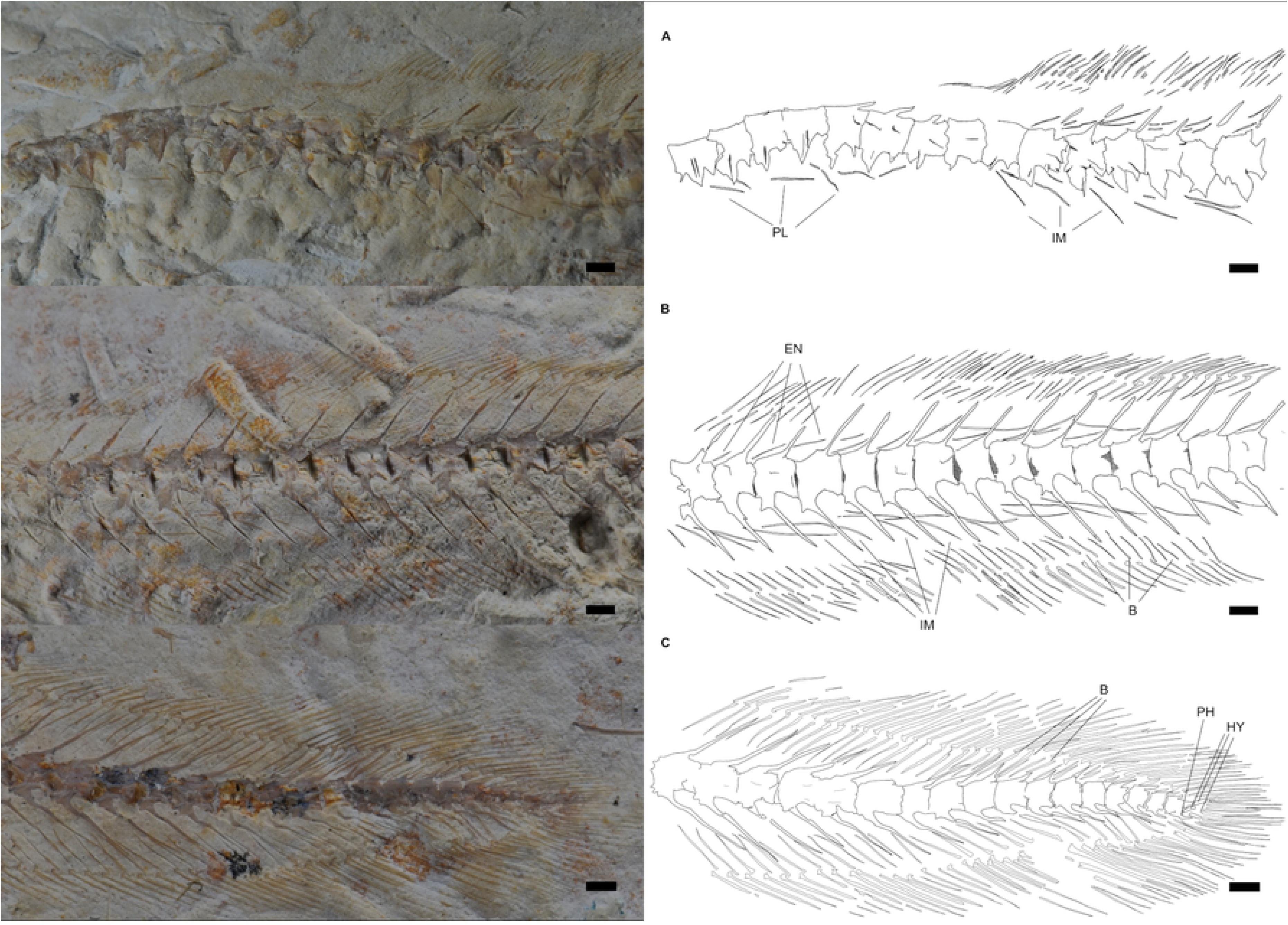
†*Mitsijt tsucul* gen. et sp. nov. paratype specimen IGM1318. Lateral view of the A) abdominal, B) caudal, and C) last vertebrae of IGM1318. Red arrows indicate yellowish patches that resemble small soft imbricated scales. Scale bar 0.5 mm.

The first dorsal pterygiophore originates in the abdominal region at the 27th vertebra, and the first anal pterygiophore is located at the 33rd vertebra. Both fins exhibit a pterygiophore concentration coefficient (number of pterygiophores per vertebra) equal to or higher than three.

Ribs (**pl**) are present in the abdominal region, beginning at the 15th vertebra. Unlike the intramuscular bones, ribs are distributed one per vertebra. The epineural (**en**) and epipleural (**im**) series begin at the dorsal and anal fins, respectively, with at least two elements per vertebra. These series consist of single, non-bifurcated bones.

### Caudal fin

As in other Anguilliformes, the dorsal and anal fins are confluent with the caudal fin. The hypural complex is formed by fused ural centra (U2+U1), with dorsal hypurals (**hy**III-IV) remaining free. In contrast, ventral hypurals (I-II) are fused into a plate and attached to the ural centra, forming a “pseudoestylar complex”. The complex also includes a hook-shaped parhypural (**ph**) (Fig. 11).

**Figure 11.**
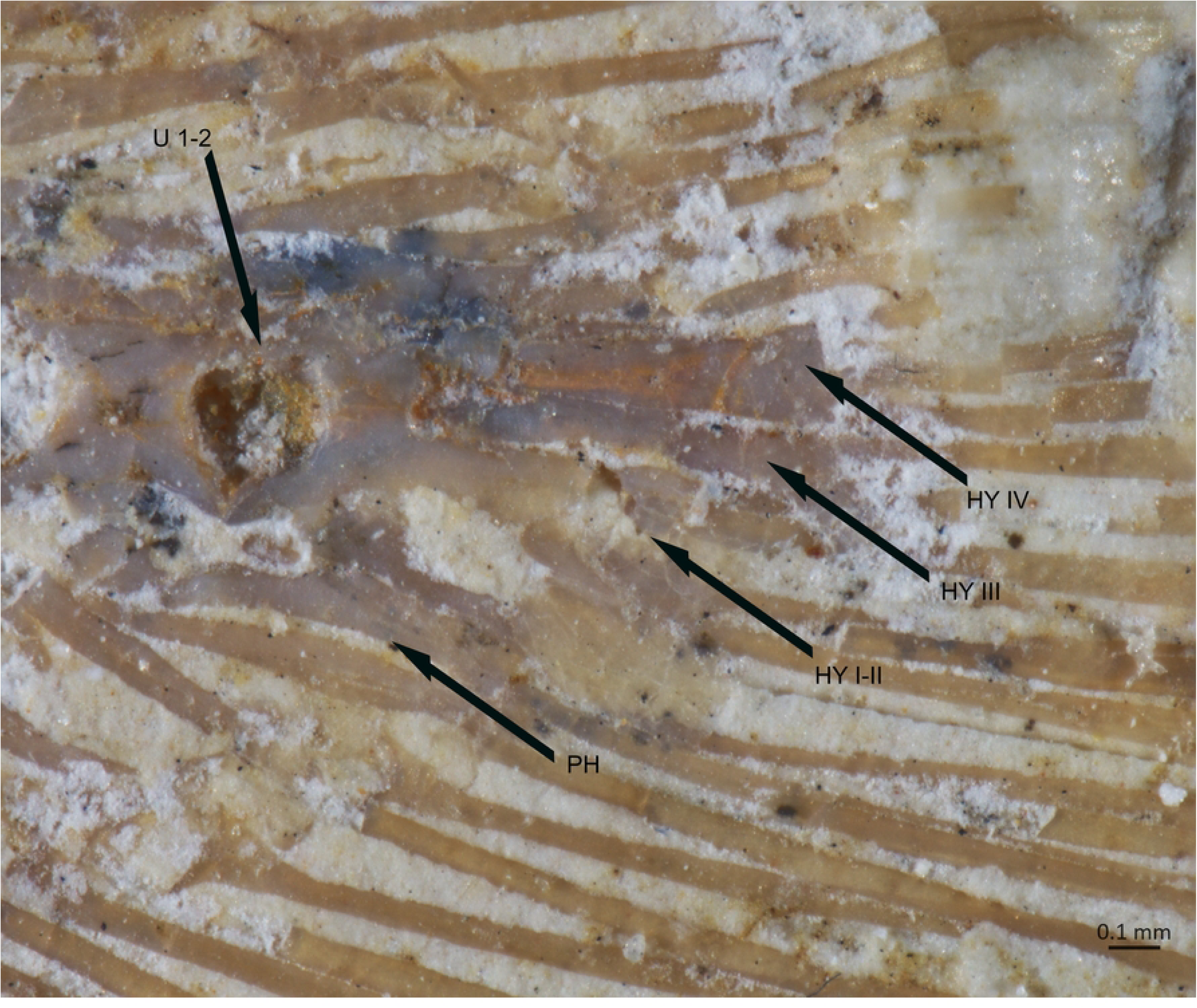
†*Mitsijt tsucul* gen. et sp. nov. paratype specimen IGM1318. Lateral view IGM1318. Scale bar 0.5.

### Phylogenetic analysis

#### Morphological phylogeny

The parsimony analysis of the complete matrix recovered 38 MPTs of 787 steps for the complete matrix and 3 MPTs of 650 for the matrix excluding Cenozoic fossils (Fig. 12). Both analyses consistently recovered: (a) a monophyletic Elopomorpha, (b) a monophyletic Anguilliformes, and (c) *P. palau* as the sister taxon of †*M. tsucul*.

**Figure 12.**
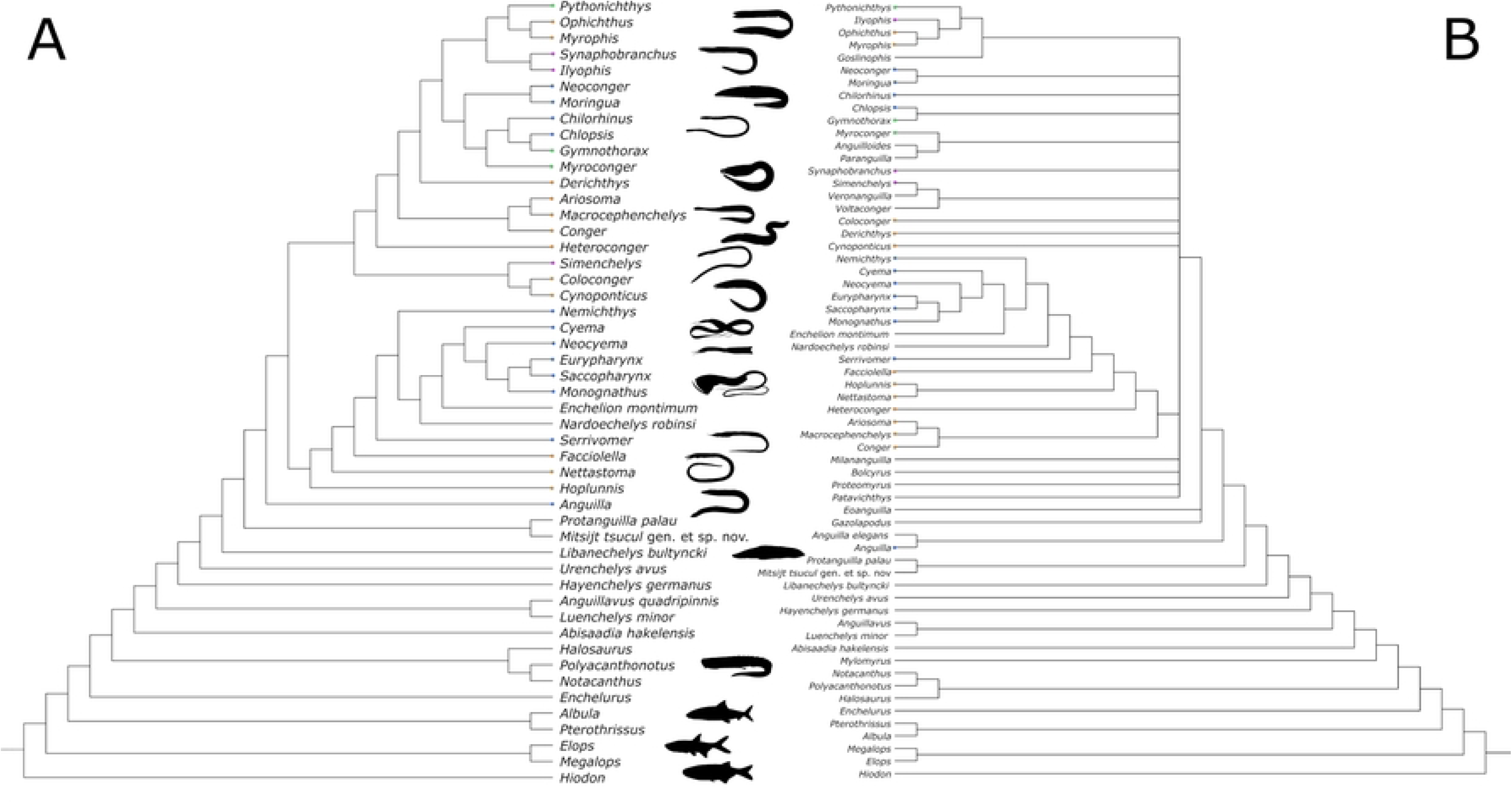
Morphological phylogeny of Anguilliformes. Semistrict consensus of the A) Cenozoic excluded morphological matrix (49 taxa, 647 steps, CI=0.329, RI= 0.728) and B) complete matrix (62 taxa, 647 steps, CI=0.329, RI= 0.728). Values on the nodes represent consensus values, dots in the terminals are coded according to the suborder of the species (purple to Synaphobranchoidei, blue to Anguilloidei, green to Muraenoidei and orange to Congroidei).

Anguilliformes is supported by the reduction of the supraoccipital (character 1, state 1), the fusion of the vomer with the ethmoids (16, 1), a double condylar articulation of the hyomandibular with the neurocranium (28, 1), the absence of the palatine (35, 2), posterior end of maxilla tapered and reaching the coronoid process (43, 1), the fusion of the post-dentary bones into an angular-articular-retroarticular complex (45, 3), caudally elongated opercular bones (50, 1), the presence of an anterodorsal branch in the subopercle (53, 1), the lack of toothed plates in the basibranchial/basihyal (57, 1), gill arches that are free from the neurocranium and posteriorly displaced (64, 1), the absence of gill rakers (65, 1), bony components of the pectoral girdle separated by cartilage (79, 1), the absence of a pectoral spine (81, 0), the first dorsal pterygiophore strongly pushed forward (89, 1), a ratio of 2.3 pterygiophores per vertebra (86, 1), discontinuity between the dorsal and caudal fins at PU4/5 (91, 1) and between the anal and caudal fins at PU5/6 (92, 1), the presence of epineural bones that are not bifurcated (117, 1), and the absence of scales (122, 2).

The crown group comprises all living eels and their related fossils, except for †*M. frangens*, which is recovered as the most basal anguilliform, and Cretaceous eels (†*N. robinsi* and †*E. montimum*), both of which are recovered as members of the Saccopharyngoidei. However, due to the lack of resolution in the affinities of Cenozoic fossils, we report synapomorphies mapped in the semi-strict consensus tree of the MPTs recovered from the Cenozoic-excluded matrix. The crown group is supported by the exact continuity of the dorsal and caudal fins (91, 3), which have fewer than eight rays per lobe (104, 0), and the fusion of PU1+U1+U2 into a pseudurostylar complex (96, 2) lacking neural arches in PU1 and U1 (103, 0).

Protanguillidae is recovered as the sister group of the rest of the crown group and is supported by the participation of the pterosphenoid in the postorbital region (7, 1), the absence of the antorbital (=adnasal) bone (21, 1), and fewer than 90 vertebrae (108, 0). *P. palau* has the dorsal wing of the sphenoid (8, 0) pointing laterally and a massive posterior end of the maxilla (43, 0) as apomorphies, but †*M. mosaicus* does not present any apomorphies.

Additionally, the “modern eels” are represented by a monophyletic group, sister to Protanguillidae, which includes all eels displaying the “typical anguilliform” body plan. This clade is supported by the fusion of the anteroventral process of the hyomandibular (31, 0) and the curvature of the subopercle behind the opercle (54, 2).

The genus *Anguilla* Linnaei, 1758 [78], is recovered as the sister taxon of all “modern eels,” which are subdivided into two major clades. The first clade comprises bathypelagic and bathydemersal eels with the anteorbital region developed as a beak. This group includes a paraphyletic Nettastomatidae, Serrivomeridae, and Nemichthyidae (members of Congroidei sensu Robins [79]) and “Saccopharyngiformes” (sensu Böhlke [60]). It is characterized by the progressive loss of opercular bones and elements of the pectoral girdle.

The sister clade of these deep-sea eels includes the rest of the extant families. The largest and best-represented families (Synaphobranchidae, Congridae, and Chlopsidae) are recovered as paraphyletic. Only Ophichthidae and Moringuidae are recovered as monophyletic, while the remaining families (Muraenesocidae, Colocongridae, Derichthyidae, Heterenchelyidae, Myrocongridae, and Muraenidae) are represented by a single taxon. This clade is supported by the presence of a lateral vomerine process (35, 1), the anterior and posterior bony contact of the pterygoid arch (37, 0), and the presence of an ossified basibranchial 3 (68, 0).

This major clade consists of Congroidei and Anguilloidei eels in a polyphyletic arrangement, but Muraenoidei (sensu Robins [79]) is recovered as monophyletic, composed of Myrocongridae as the sister taxon to Chlopsidae and Muraenidae. This clade is supported by the loss of basibranchial 1 (66, 1) and ribs (119, 2). The relationship between Chlopsidae and Muraenidae is supported by the presence of bony protection of the anterior supraorbital branch in the frontal canal (23, 0), the reduction of the preopercle (51, 1) and interopercle (52, 1), the loss of hypobranchial 3 (69, 2), the reduction or loss of bony elements in the pectoral girdle (79, 3), and the absence of pectoral fins (80, 2).

### Total evidence phylogeny

Our analysis was congruent with the morphologic analysis in the recovery of (a) a monophyletic Elopomorpha, (b) a monophyletic Anguilliformes, and (c) *P. palau* as the sister taxon of †*M. tsucul* (Fig. 13).

**Figure 13.**
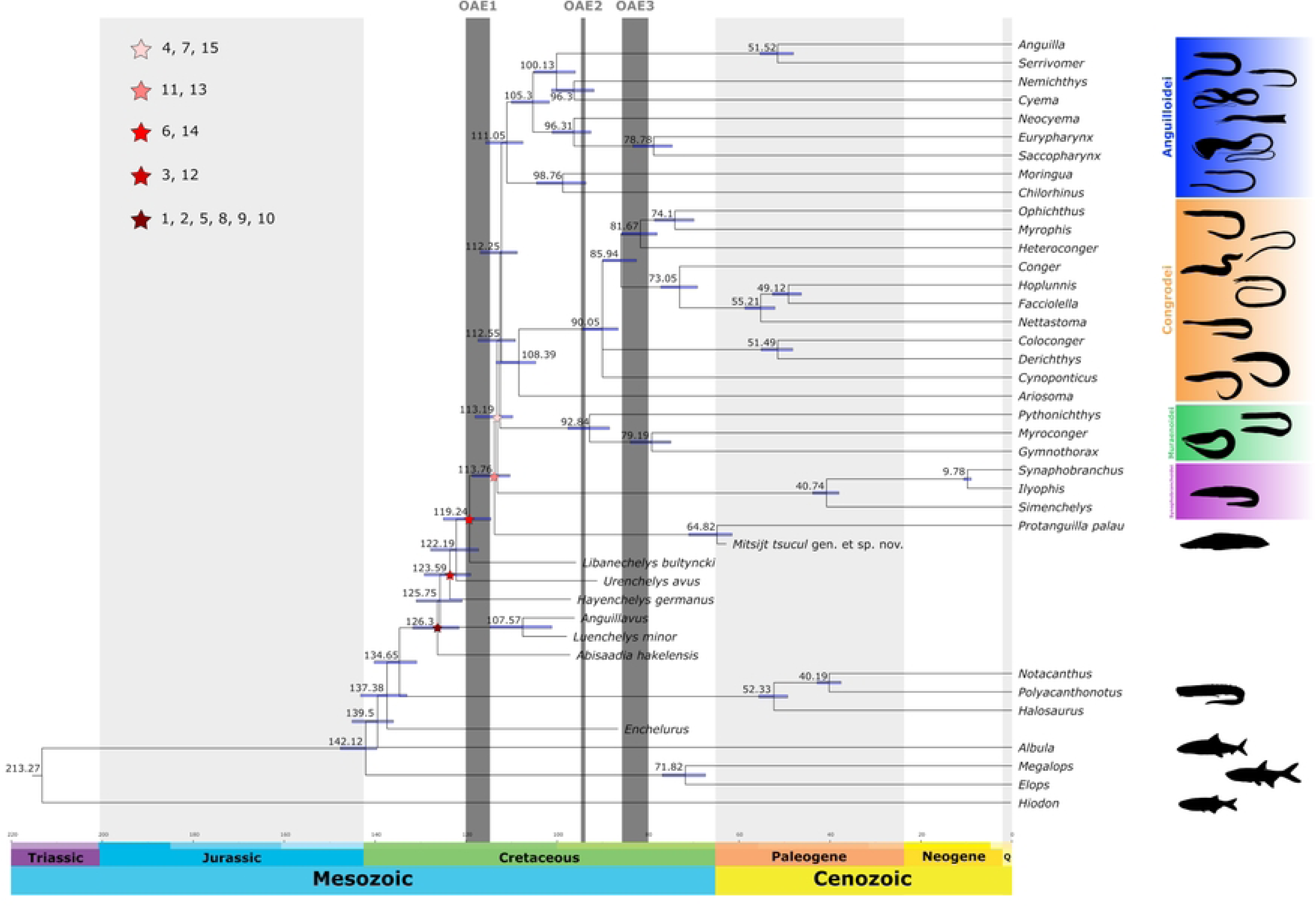
Tip dating time divergence estimation of Anguilliformes based on morphology and mitochondrial genes. Phylogenetic relationships of Anguilliformes from the total evidence matrix with topological constraints on Cretaceous eels based on the parsimony analysis. Synapomorphies proposed of Anguilliformes by Robins [77] and coded in the Belouze’s matrix, *Anguilla* represents the modern eels. We notice that the disposition of pterotic reaching or exceeding the sphenoid (1), fusion of vomer in the ethmovomer (2), absence of palatine (5), fusion of anguloretroarticular (8), the posterocaudally elongated opercular bones (9) and the posterior displacement of branchial arches (10) are synapomorphies of pan-anguilliforms but absence of lateral ethmoid (3), fusion of endopterygoid and ectopterygoid in a single arc (6), absence of posttemporals (11) and pelvic girdle (12) and confluence of unpaired fins (13, 14) are acquired during the Late Cretaceous. Synapomophies of the modern eels absent in stem eels are absence of scales (15) (excepting *Anguilla*), fusion of symplectic (4) and premaxilla (7). These two last characters are also present in †Libanechelyidae but no in Protanguillidae.

Anguilliformes is supported by the same synapomorphies of previous morphological analysis, the crown-Anguilliformes are supported by the same characters (91, 96, 103) except for the presence of unfused epineural bones to neural arches (118, 1). The modern eels are supported by the supraoccipital with a straight posterior edge (2, 1), moderately elongated anteorbital region (15, 2), an interdigitated suture of hyomandibular and quadrate (34, 2), the absence of neurospine in PU2 (98, 2) and upper hypurals fused in a plate (100, 1).

The origin of Elopomorpha was estimated to be in the Early Cretaceous (142.12 Ma; 95% Highest Posterior Density: 147.66-139.58 Ma). The origin of crown Elopiformes is recovered during the Late Cretaceous (71.82 Ma., 95% HPD= 67.35-76.85 Ma.), while the split between Albuliformes and Anguilliformes + Notacanthiformes is recovered in the Early Cretaceous (139.5 Ma., 95% HPD= 136.02-145.08 Ma.). The split of Notacanthiformes and Anguilliformes occurred in the Early Cretaceous (134.65 Ma., 95% HDP= 130.84-140.22 Ma), but the origin of crown Notacanthiformes is recovered in the Eocene (52.33 Ma., 95% HDP= 49.3-55.65). The origin of Anguilliformes recovered in the Early Cretaceous (126.3 Ma., 95% HDP= 121.57-131.71 Ma.) but the crown group appeared later in the Early Cretaceous (113.76 Ma., 95% HDP= 110.37-118.64 Ma) and the modern eels diversified shortly thereafter in the Early Cretaceous (113.19 Ma., 95% HDP= 109.76-118.01.). The split between †*M. tsucul* and *P. palau* are recovered in the early Paleocene (64.82 Ma., 95% HDP = 61.51-71.01 Ma.).

Finally, the SS test was inconclusive: the total evidence matrix obtained a better marginal likelihood score (-226147.69) in the unconstrained analysis (Protanguillidae nested in Synaphobranchidae) than the constrained Protanguillidae (-226161.81) and †Libanechelyidae (-226161.88) hypotheses. However, morphological data tests provided better support for the †Libanechelyidae hypothesis (-2613.92) than for the Protanguillidae hypothesis (-2650.53) and the unconstrained topology (-2658.65).

## Discussion

### The phylogenetic position of **†***Mitsijt tsucul* gen. et sp. nov. in Protanguillidae

The inclusion of †*M. tsucul* gen. et sp. nov. within Anguilliformes is supported by key features such as the fusion of the mesethmoid and vomer (16, 1), as well as the angular, articular, and retroarticular bones (45, 3). Additional traits include the arrangement of the opercular series (50, 1), the absence of a pelvic girdle (85, 2), and confluent unpaired fins (91, 3; 92, 3) [60,76]. However, †*M. tsucul* retains some plesiomorphic characters absent in modern eels but shared with Cretaceous fossils and *P. palau*. These include an autogenous premaxilla (39, 0), free symplectic (33, 0) and metapterygoid bones, and unfused dorsal hypurals (110, 0; 102, 0). Notably, some specimens exhibit small and imbricated scales, and specimen IGM1307 possesses a structure in a congruent position to the gill rakers of *P. palau*. While its identity remains unclear, the existence of both features in †*M. tsucul* is plausible, as seen in extant species.

Unfortunately, the muscular characters of the gill arches [37] and cephalic structures [38], which support the basal position of the lineage in *P. palau*, are unknown in †*M. tsucul*. Nevertheless, the osteological traits are sufficient to assign this new species to the peculiar family Protanguillidae.

Despite significant discordance among the morphological and total evidence phylogenetic analyses performed in this study, all consistently recover †*M. tsucul* as the sister species to *P. palau* with strong support. This includes a bootstrap value of 60 in the complete matrix, 61 in the matrix excluding Cenozoic fossils, and a posterior probability of 1 in the total evidence analysis.

### Origin and early evolution of Anguilliformes

Our divergence time estimation places the origin of Elopomorpha in the Late Jurassic, consistent with the oldest elopomorph fossils and most previous estimates (Fig. 13). The origin of Anguilliformes is recovered in the Early Cretaceous, also in agreement with previous estimates [35,80–83], but creates a gap of approximately 20 million years with the oldest known anguilliform fossils.

The oldest fossil record of true eels dates to the Cenomanian of Lebanon [11,13] and includes eels with plesiomorphic traits [8–10] which have caused doubt about their relationship with living anguilliforms [79]. However, our results support the interpretation of these forms as stem-group anguilliforms [7] and include highly modified eels with similar morphology to living abyssal eels [16,84], indicating a high specialization in the early evolution of the group.

The phylogenetic position of protanguillids relative to both Cretaceous and extant anguilliforms has remained uncertain due to their distinctive morphology [6,37,38]. Previous tests of alternative topologies have relied exclusively on molecular data [6,25], and the affinities of †*L. bultyncki*, which also exhibits derived features comparable to protanguillids [14], had not been evaluated.

Our total-evidence analyses were inconclusive with respect to the placement of †*L. bultyncki*, as marginal likelihood values did not differ significantly between constrained hypotheses. This result may reflect both the limited basis for coding the species †*L. bultyncki* (restricted to its original description) and the difficulty of integrating fragmentary fossil taxa into combined datasets.

Nonetheless, the combined molecular and morphological evidence supports recognition of Protanguillidae as a surviving lineage of Cretaceous anguilliforms rather than as part of the modern eel radiation. The sister-group relationships of crown anguilliforms remain unresolved, underscoring the need for additional fossil material and expanded morphological sampling to clarify early divergences within the group.

Our morphological phylogeny recovers Protanguillidae as sister taxa of the modern eels, and we set this topology as prior in our TD analysis. This decision is based on the following characters: †*L. bultyncki* retains an endopterygoid, ectopterygoid, palatine, posttemporal bones, and a discontinuity between dorsal and caudal fin [14], characters present in its derived condition of Protanguillidae. In turn, protanguillids present autogenous premaxilla and symplectic, gill rakers, scales and the pterosphenoids as part of the posterior orbit margin. However, †*L. bultyncki* is known from a single specimen preserved in dorsal view, which limits our ability to confirm certain key features, such as the partial fusion of the premaxilla and details of the orbital region. Additionally, scales and gill rakers are rarely preserved in such fossils, and more material is needed to clarify their affinities.

Before the discovery of protangulids the identification of the most basal lineage of true eels was problematic. Based on anatomy [37,85] and osteology [79,86] Anguillidae, Congridae, and Synaphobranchidae have been proposed as the earliest-diverging true eels. Molecular phylogenies have recognized Synaphobranchoidei as the oldest lineage of modern eels, but the divergence of the living synaphobranchoids is estimated to have occurred in the Eocene (40.74 Ma.). Synaphobranchoids retain some plesiomorphic traits as a “cartilaginous symplectic”, small and imbricated scales arranged in a “basket-weave” pattern, and up to five free hypurals in some *Synaphobranchus* Johnson, 1862 species [87–92]. This combination of plesiomorphic and modern traits can be considered as evidence of the affinities of Protanguillidae to Synaphobranchoidei, as suggested by the ss test of the total evidence matrix.

Finally, the Danian outcrops in Palenque, Mexico, have yielded some of the oldest known fossils of reef fishes as trumpetfishes [49,93], damselfishes [47,94] and seabasses [46,95]. Previously, the oldest fossil record for many reef fishes was the Eocene Monte Bolca locality in Italy [18,20,96]. The discovery of these Paleocene fossils and the use of TD analysis suggests the apparition of these lineages near the Cretaceous-Paleogene boundary [41]. The only living protanguilid, *P. palau*, was discovered in a reef cave of the Republic of Palau [6] and our divergence time estimation suggests the divergence between *P. palau* and †*M. tsucul* during the early Danian (64.82 Ma.), in accordance with the estimated divergence times for the other reef fishes of Palenque.

### The gradual acquisition of modern eel traits

The species of living true eels have a long and problematic taxonomic history, due to the widespread anguilliform body plan among several fish lineages. However, the delimitation of true eels as Apodes -later formalized as Anguilliformes- has been made possible by key osteological traits [75,76,78,79,97,98].

Here we defined modern eels as the sister group of Protanguillidae, described as the most inclusive clade that contains *Synaphobranchus oregoni* Castle, 1960 [52] and *Eurypharynx pelecanoides* Vaillant, 1882 [99] but not *Protanguilla palau* Johnson, Ida & Sakaue, 2012 [6]. Nevertheless, we have found that the osteological traits used to delimit Anguilliformes emerged gradually in the Late Cretaceous within the stem group, and only a few are synapomorphies of the crown group and the modern eels.

We recovered congruent relationship of the four suborders of modern eels (Synaphobranchoidei, Anguilloidei, Muraenoidei and Congroidei) congruent with previous studies [6,25,39] except for the position of Chlopsidae. In our analysis, Chlopsidae is recovered as the sister group of Moringuidae within Anguilloidei, whereas previously studies recovered it as a congroid [6,39,65,100], as muraenoid [25] or as an independent lineage [1,33]. The use of TD method allowed us to identify a rapid diversification of the crown group and modern eels during the Early Cretaceous after the Oceanic Anoxic Event I (Fig. 14), a pattern also observed in acanthomorphs [83,101–103] and elasmobranchs [104–106]. However, unlike other fish lineages, no body fossils of modern eels from the Cretaceous have been described.

Nevertheless, the otolith fossil record provides the existence of modern eels during the Late Cretaceous with the presence of congrids, anguillids and protanguillids [28–30,107]. Additionally, numerous modern eels are recognized in the otolith record during the Cenozoic as anguillids, ophicthids, congrids heterenchelyds, moringuids, nettastomatids, muraenesocids and protanguillids during the Paleogene [26,27,30,108–111] and Neogene [112–114]. Despite this extensive record of modern eels otoliths and its general the congruence with our divergence time estimation, we are sceptic about the validity of the taxonomic assignment of otoliths to modern genera and families.

The oldest known fossils that can be reliably assigned to modern eels come from the Eocene Monte Bolca locality [18,20]. These fossils exhibit all the modern eels traits, but their phylogenetic relationships remain poorly resolved. Some of the taxa present has been classified in a living family, as †*Whitapodus breviculus* Blot, 1980 [20], which he assigned to Xenocongridae (=Chlopsidae) based on opercular reduction and the absence of pectoral fins; †*Goslinophis acuticaudus* Blot, 1980 [20], which he classified as an ophichthid based on the inclination of the suspensorium and the position of the dorsal and anal fins; and †*Paranguilla tigrina* (Agasizz, 1839) [115], related to muraenids based on the inclination of the suspensorium, the lack of neural spines, and the reduction of the pectoral girdle [20,115]. However, more detailed descriptions and systematic study are required.

Other Cenozoic eels have been assigned to extant families based on exhaustive osteological comparisons, such as †*Bohlkea elongata* Taverne & Nolf, 1978 [116] (Chlopsidae) and †*Nettastoma belgica* Taverne & Nolf, 1978 (Nettastomatidae) [116]. However, we do not fully agree with the assignment of †*N. belgica* to this genus due to the resemblance of its anteriormost ethmovomer and dentary segments to those of other genera, although its affinities with nettastomatids are apparent. Similarly, †*Echelus branchialis* (Woodward, 1901) [13] has been compared to other species of the same genus [117], †*Deutschenchelys micklichi* (Prokofiev, 2012) [23] (Moringuidae) is characterized by burrowing adaptations resembling those of *Neoconger* Girard, 1858 [118], and *Moringua* Gray, 1831 [23,118,119] and †*Serrivomer glehni* Nazarkin, 2023 [24] distinguished by clear diagnostic characters present in the living serrivomerids [24].

Nevertheless, our results suggest that the assignment of fossil taxa to Congridae should be approached with caution. Heterocongrinae is a specialized group of burrowing eels closely related to Bathymyrinae. The assignment of †*Pavelichthys daniltshenkoi* Prokofiev, 2007, to this group, based on the disposition of the abdominal and caudal vertebrae and cranial architecture, seems reasonable [22]. However, the classification of †*Smithconger treldeensis* (Schwarzhans, 2007) [120] within Congrinae [21] is problematic due to the high morphological diversity of congrids and the polyphyly of the group recovered in this study and other molecular phylogenies [25].

## Conclusions

Here, we describe †*Mitsijt tsucul* gen. et sp. nov. a new anguilliform fish belonging to the family Protanguillidae. This fossil represents the first unambiguous record of a living lineage of Anguilliformes in the fossil record and the second known genera of Protanguillidae. Additionally, we present the most extensive phylogenetic analysis of combined data to date, including Cretaceous taxa, to test the positions of †Libanenchelyidae and Protanguillidae as sister taxa to all extant eels.

We show that the early evolutionary history of eels during the Cretaceous remains largely unresolved, mainly due to the scarcity of fossils that can be confidently assigned to the crown group. Our results reveal a significant diversification among major lineages during the Late Cretaceous and indicate that many of the traditional synapomorphies defining the order were gradually acquired along the stem lineage.

Additionally, we suggest an exhaustive taxonomic revision of the oldest Cenozoic fossils to clarify the evolutionary history of true eels after the K/Pg boundary to shed light on the survival of primitive lineages to the present day.

## Supporting information

**S1 PDF.** Supporting information. A pdf containing the list of characters coded in the morphological matrix and a review of the codification of †*M. tsucul*, the character mapping of the strict consensus tree of the Cenozoic excluded matrix and the Total Evidence tree and the morphological matrix.

(PDF)

**S2 File. Discrete morphological matrix**

(NEX)

**S3 File. Total evidence matrix**

(NEX)

## Acknowledgments

This manuscript is requirement for the first author (SHB) to to obtain the degree of Master in Biological Sciences (Systematics) within the *Posgrado en Ciencias Biológicas* at *Universidad Nacional Autónoma de México*. I (SHB) am grateful to CONAHCYT (now SECIHTI) for funding and for the support of this research. Thanks to M.Sc. Christian Lambarri Martínez and Dr. Wilfredo Matamoros Ortega for their invaluable support and guidance, to Dra. Marisol Montellano Ballesteros and Dr. Luis Fernando del Moral for their participation during the development of this project. Thanks to M.Sc. Violeta Amparo Romero Mayen for cataloging and storing the specimens in the type collection. We are grateful with M.Sc. Camila Alcantara from *Laboratorio de Microscopía y Fotografía de la Biodiversidad* (II), IB-UNAM, for her advice and assistance in obtaining microphotographs and Dr. Joseph Anton Moreno Bedmar for his assistance in the photographing of the specimens described.

## Author contributions

**Conceptualization:** Sebastián Huacuja-Barraza, Kleyton M Cantalice.

**Data curation:** Sebastián Huacuja-Barraza, Kleyton M Cantalice.

**Funding acquisition:** Kleyton M Cantalice.

**Investigation:** Sebastián Huacuja-Barraza.

**Methodology:** Sebastián Huacuja-Barraza.

**Project administration:** Sebastián Huacuja-Barraza, Kleyton M Cantalice.

**Resources:** Kleyton M Cantalice.

**Validation:** Kleyton M Cantalice.

**Visualization:** Sebastián Huacuja-Barraza, Kleyton M Cantalice.

**Writing – original draft:** Sebastián Huacuja-Barraza.

**Writing – review & editing:** Sebastián Huacuja-Barraza, Kleyton M Cantalice

## References

1. Near TJ, Thacker CE. Phylogenetic Classification of Living and Fossil Ray-Finned Fishes (Actinopterygii). Bull Peabody Mus Nat Hist. 2024;65: 3–302. doi:10.3374/014.065.0101

2. Fricke R, Eschmeyer WN, Van der Laan R. Eschmeyer’s Catalog of Fishes: Genera, species, references. In: Eschmeyer’s Catalog of Fishes [Internet]. 2025. Available: http://researcharchive.calacademy.org/research/ichthyology/catalog/fishcatmain.asp

3. Nelson J, Grande T, Wilson MVH. Fishes of the world. 5th ed. John Wiley & Sons; 2016.

4. Helfman GS, Collette BB, Facey DE, Bowen BW. The diversity of fishes: biology, evolution, and ecology. 2nd ed. Atlantic. Wiley-Blackwell; 2009. doi:10.1007/978-1-4615-2664-3_1

5. Robins CR. The Phylogenetic Relationships of the Anguilliform Fishes. In: Bohlke E, editor. Fishes of the Western North Atlantic. Sears Foundation for Marine Research; 1989.

6. Johnson GD, Ida H, Sakaue J, Sado T, Asahida T, Miya M. A “living fossil” eel (Anguilliformes: Protanguillidae, fam. nov.) from an undersea cave in Palau. Proceedings of the Royal Society B: Biological Sciences. 2012;279: 934–943. doi:10.1098/rspb.2011.1289

7. Belouze A. Compréhension morphologique et phylogénétique des taxons actuels et fossiles rapportés aux Anguilliformes (Poissons», Téléostéens). Travaux et Documents des Laboratoires de Géologie de Lyon. Université Claude Bernard Lyon 1. 2002. Available: https://www.persee.fr/doc/geoly_0750-6635_2002_mon_158_1

8. Belouze A, Gayet M, Atallah C. The first Anguilliformes: I. Revision of the Cenomanian genera Anguillavus HAY, 1903 and Luenchelys nov. gen. Geobios. 2003;36: 241–273. doi:10.1016/S0016-6995(03)00029-9

9. Belouze A, Gayet M, Atallah C. The first Anguilliformes: II. Paraphyly of the genus Urenchelys WOODWARD, 1900 and phylogenetic relationships. Geobios. 2003;36: 351–378. doi:10.1016/S0016-6995(03)00036-6

10. Belouze A, Gayet M. Premiere observation de la presence simultanee d’une ceinture pectorale complete et de nageoires pelviennes chez des Anguilliformes (Teleostei) du Cretace du Liban. Comptes Rendus de l’Academie de Sciences - Serie IIa: Sciences de la Terre et des Planetes. 1999;329: 683–688. doi:10.1016/S1251-8050(00)87646-6

11. Hay O. On A Collection Of Upper Cretaceous Fishes From Mount Lebanon, Syria, With Descriptions Of Four New Genera And Nineteen New Species. Bull Am Mus Nat Hist. 1903;19.

12. Woodward AS. XXLIV. —Evidence of an extinct eel (Urenchelys anglicus, sp. n) from the English Chalk. J Nat Hist. 1900;5: 321–323. doi:10.1080/00222930008678294

13. Woodward AS. Catalogue of the fossil fishes. British Museum (Natural History). 1901;4.

14. Taverne L. Libanechelys bultyncki gen. et sp. nov., une nouvelle anguille primitive (Teleostei, Anguilliformes) du Cénomanien marin du Liban. Sciences de la Terre. 2004; 73–87.

15. Taverne L. Les poissons crétacés de Nardo. 13° Nardoechels robinsi gen. et sp. nov., la plus ancienne anguille de type moderne connue par des élélments squelettiques (Teleostei, Anguilliformes, Ophichthidae). Boll Soc Paleontol Ital. 2002;26: 25–31.

16. Taverne L, Capasso L. Les poissons crétacés de Nardò. 36°. Compléments à l’étude de Nardoechelys robinsi Taverne, 2002 (Teleostei, Anguilliformes). Geologia Paleontologia Preistoria. 2014.

17. Woodward AS. VI. —On a Fossil Sole and a Fossil Eel from the Eocene of Egypt. Geol Mag. 1910;7: 402–405. doi:10.1017/S0016756800135186

18. Bannikov AF. The systematic composition of the Eocene actinopterygian fish fauna from Monte Bolca, northern Italy, as known to date. Miscellanea Paleontologica. 2014;12: 22–34.

19. Carnevale G, Bannikov AF, Marramà G, Tyler JC, Zorzin R. The Pesciara-Monte Postale Fossil-Lagerstätte: 2. Fishes and other vertebrates. Rendiconti della Società Paleontologica Italiana. 2014;4: 37–63.

20. Blot J. La faune ichtliyologique des gisements du Monte Bolca (Province de Vérone, Italie). Catalogue systématique présentant l’état actuel des recherches concernant cette faune. Bulletin du Museum National d’Histoire Naturelle. 1980; 339–396.

21. Carnevale G, Schwarzhans W, Schrøder AE, Lindow BEK. An Eocene conger eel (Teleostei, Anguilliformes) from the Lillebælt Clay Formation, Denmark. Bulletin of the Geological Society of Denmark. 2022;70: 53–67. doi:10.37570/bgsd-2022-70-05-rev

22. Prokofiev AM. A redescription and relationships of the congrid eel Pavelichthys daniltshenkoi (Anguilliformes: Congridae) from the lower Oligocene of Northern Caucasus. J Ichthyol. 2007;47: 335–340. doi:10.1134/S0032945207050013

23. Prokofiev AM. Oligocene eel from the Frauenweiler site (Germany). J Ichthyol. 2012;52: 11–18. doi:10.1134/S0032945212010109

24. Nazarkin M V. A saw-toothed eel †Serrivomer glehni sp. nov. from the Miocene of Sakhalin Island, north-western Pacific. J Vertebr Paleontol. 2023. doi:10.1080/02724634.2023.2261505

25. Santini F, Kong X, Sorenson L, Carnevale G, Mehta RS, Alfaro ME. A multi-locus molecular timescale for the origin and diversification of eels (Order: Anguilliformes). Mol Phylogenet Evol. 2013;69: 884–894. doi:10.1016/j.ympev.2013.06.016

26. Schwarzhans W. Fish otoliths from the Paleocene of Bavaria (Kressenberg) and Austria (Kroisbach and Oiching-Graben). Palaeo Ichthyologica. 2012.

27. Schwarzhans W. Fish otoliths from the Paleocene of Denmark. Geological Survey of Denmark and Greenland Bulletin. 2003. doi:10.34194/geusb.v2.4696

28. Stringer G, Schwarzhans W. Upper Cretaceous teleostean otoliths from the Severn Formation (Maastrichtian) of Maryland, USA, with an unusual occurrence of Siluriformes and Beryciformes and the oldest Atlantic coast Gadiformes. Cretac Res. 2021;125: 104867. doi:10.1016/j.cretres.2021.104867

29. Schwarzhans W, Jagt JWM. Silicified otoliths from the Maastrichtian type area (Netherlands, Belgium) document early gadiform and perciform fishes during the Late Cretaceous, prior to the K/Pg boundary extinction event. Cretac Res. 2021;127: 104921. doi:10.1016/j.cretres.2021.104921

30. Schwarzhans W, Stringer GL. Fish otoliths from the Late Maastrichtian Kemp clay (Texas, USA) and the Early Danian Clayton Formation (Arkansas, USA) and an assessment of extinction and survival of teleost lineages across the K-Pg boundary based on otoliths. Rivista Italiana di Paleontologia e Stratigrafia. 2020;126: 395–446.

31. Forey PL, Littlewood DTJ, P. Ritchie, Meyer A. Interrelationships of Elopomorph Fishes. In: Stiassny M, Parenti LR, Johnson GD, editors. Interrelationships of Fishes. San Diego: Academic Press; 1996. pp. 174–191.

32. Arratia G. Basal Teleost and Teleostean Phylogeny. Palaeo Ichthyologica. 1997;7.

33. Wang CH, Kuo CH, Mok HK, Lee SC. Molecular phylogeny of elopomorph fishes inferred from mitochondrial 125 ribosomal RNA sequences. Zool Scr. 2003;32: 231–241. doi:10.1046/j.1463-6409.2003.00114.x

34. Filleul A, Lavoué S. Basal teleosts and the question of elopomorph monophyly. Morphological and molecular approaches. Académie des sciences/Éditions scientifiques et médicales. 2001;324: 393–399. doi:10.1016/S0764-4469(00)01302-0

35. Dornburg A, Friedman M, Near TJ. Phylogenetic analysis of molecular and morphological data highlights uncertainty in the relationships of fossil and living species of Elopomorpha (Actinopterygii: Teleostei). Mol Phylogenet Evol. 2015;89: 205–218. doi:10.1016/j.ympev.2015.04.004

36. Inoue JG, Miya M, Tsukamoto K, Nishida M. Mitogenomic evidence for the monophyly of elopomorph fishes (Teleostei) and the evolutionary origin of the leptocephalus larva. Mol Phylogenet Evol. 2004;32: 274–286. doi:10.1016/j.ympev.2003.11.009

37. Springer VG, Johnson GD. The Gill-Arch Musculature of Protanguilla, the Morphologically Most Primitive Eel (Teleostei: Anguilliformes), Compared with That of Other Putatively Primitive Extant Eels and Other Elopomorphs. Copeia. 2015;103: 595–620. doi:10.1643/CI-14-152

38. Espíndola VC, Johnson GD, De Pinna MCC. Facial and opercular muscles in the Anguilliformes (Elopomorpha: Teleostei): Comparative anatomy and phylogenetic implications for the basal position of Protanguilla. J Morphol. 2023;284: 1–13. doi:10.1002/jmor.21556

39. Tang KL, Fielitz C. Phylogeny of moray eels (Anguilliformes: Muraenidae), with a revised classification of true eels (Teleostei: Elopomorpha: Anguilliformes). Mitochondrial DNA. 2013;24: 55–66. doi:10.3109/19401736.2012.710226

40. Chen J-N, Andrés López J, Lavoué S, Miya M, Chen W-J. Phylogeny of the Elopomorpha (Teleostei): Evidence from six nuclear and mitochondrial markers. Mol Phylogenet Evol. 2014 [cited 8 Aug 2023]. doi:10.1016/j.ympev.2013.09.002

41. Cantalice KM, Alvarado-Ortega J, Bellwood DR, Siqueira AC. Rising from the Ashes: The Biogeographic Origins of Modern Coral Reef Fishes. Bioscience. 2022;72: 769–777. doi:10.1093/biosci/biac045

42. Alvarado-Ortega J, Cuevas-García M, Melgarejo-Damián MDP, Cantalice KM, Alaniz-Galvan A, Solano-Templos G, et al. Paleocene fishes from palenque, Chiapas, southeastern Mexico. Palaeontol Electronica. 2015;18: 1–22. doi:10.26879/536

43. Alvarado-Ortega J, Cuevas-García M, Cantalice KM. The fossil fishes of the archaeological site of Palenque, Chiapas, southeastern Mexico. J Archaeol Sci Rep. 2018;17: 462–476. doi:10.1016/j.jasrep.2017.11.029

44. Cuevas-García M, Alvarado-Ortega J. Estudio arqueológico y paleontológico de los fósiles marinos que proceden del sitio de Palenque, Chiapas, Informe de la primera temporada de campo. 2009.

45. Guadarrama A, Cantalice KM. Two contemporaneous morphs of fossil Chanos Lacepède, 1803 (Gonorynchiformes, Chanidae) from Paleocene (Danian) outcrops near Palenque (Mexico) revealed by geometric morphometrics indicate conservatism in milkfishes after the K / Pg boundary. PLoS One. 2025;1803. doi:10.1371/journal.pone.0313912

46. Cantalice KM, Alvarado-Ortega J, Alaniz-Galvan A. Paleoserranus lakamhae gen. et sp. nov., a Paleocene seabass (Perciformes: Serranidae) from Palenque, Chiapas, southeastern Mexico. J South Am Earth Sci. 2018;83: 137–146. doi:10.1016/j.jsames.2018.01.010

47. Cantalice KM, Alvarado-Ortega J, Bellwood DR. †Chaychanus gonzalezorum gen. et sp. nov.: A damselfish fossil (Percomorphaceae; Pomacentridae), from the Early Paleocene outcrop of Chiapas, Southeastern Mexico. J South Am Earth Sci. 2020;98: 102322. doi:10.1016/j.jsames.2019.102322

48. Cantalice KM, Alvarado-Ortega J. Kelemejtubus castroi, gen. et sp. nov., an ancient percomorph (Teleostei, Actinopterygii) from the Paleocene marine deposits near Palenque, Chiapas, southeastern Mexico. J Vertebr Paleontol. 2017;37. doi:10.1080/02724634.2017.1383265

49. Alvarado-Ortega J, Cantalice KM. Eekaulostomus cuevasae gen. and sp. nov., an ancient armored trumpetfish (Aulostomoidea) from Danian (Paleocene) marine deposits of Belisario Domínguez, Chiapas, southeastern Mexico. Palaeontol Electronica. 2016;19: 1–24.

50. Toombs HA, Rixon AE. The use of plastics in the” Transfer Method” of preparing fossils. The Museums Journal. 1950;50.

51. McCosker J. The osteology, classification and relationships of the eel family Ophichthidae. Proceedings of the California Academy of Sciences. San Francisco California Academy of Sciences; 1977. pp. 1–123. Available: http://biostor.org/reference/59597

52. Castle PH. Two eels of the genus Synaphobranchus from the Gulf of Mexico. Chicago: Chicago Natural History Museum; 1960.

53. Gosline WA. The Osteology and Relationships of the Echelid Eel, Kaupichthys diodonius. Pac Sci. 1950;4: 309–314.

54. Gosline WA. The osteology and classification of the ophichthid eels of the Hawaiian Islands. Pac Sci. 1951;5: 298–320.

55. Robins CH, Robins CR. The eel family Dysommidae (including the Dysomminidae and Nettodaridae), its osteology and composition, including a new genus and species. Academy of Natural Sciences. 1970;122: 293–335.

56. Nakamura K, Mushiake K. Anatomy on the Opercular System of the Eel, Anguilla japonica. Memoirs of Faculty of Fisheries Kagoshima University. 1982; 103–111.

57. De Schepper N, Adriaens D, De Kegel B. Moringua edwardsi (Moringuidae: Anguilliformes): Cranial specialization for head-first burrowing? J Morphol. 2005;266: 356–368. doi:10.1002/jmor.10383

58. Eagderi S, Adriaens D. Cephalic morphology of Pythonichthys macrurus (Heterenchelyidae: Anguilliformes): Specializations for head-first burrowing. J Morphol. 2010;271: 1053–1065. doi:10.1002/jmor.10852

59. Smith DG, Irmak E, Özen Ö. A Redescription of the Eel Panturichthys fowleri (Anguilliformes: Heterenchelyidae), with a Synopsis of the Heterenchelyidae. Copeia. 2012; 484–493. doi:10.1643/CI-11-174

60. Bohlke E, Bohlke J, Leiby M, McCosker J, Bertelsen E, Robins C, et al. Fishes of the Western North Atlantic Part Nine: Orders Anguilliformes and Saccopharyngiformes. Bohlke E, editor. Sears Foundation for Marine Research; 1989.

61. Maddison WP, Maddison DR. Mesquite: a modular system for evolutionary analysis. 2023. Available: http://www.mesquiteproject.org

62. Castro-Aguirre JL, Espinosa Pérez H, Schmitter-Soto JJ. Ictiofauna estuarino-lagunar y vicaria de México. Mexico City: Editorial Limusa; 1999.

63. Goloboff PA, Morales ME. TNT version 1.6, with a graphical interface for MacOS and Linux, including new routines in parallel. Cladistics. 2023;39: 144–153. doi:10.1111/cla.12524

64. Inoue JG, Miya M, Tsukamoto K, Nishida M. Evolution of the Deep-Sea Gulper Eel Mitochondrial Genomes: Large-Scale Gene Rearrangements Originated Within the Eels. Mol Biol Evol. 2003;20: 1917–1924. doi:10.1093/molbev/msg206

65. Poulsen JY, Miller MJ, Sado T, Hanel R, Tsukamoto K, Miya M. Resolving deep-sea pelagic saccopharyngiform eel mysteries: Identification of Neocyema and Monognathidae leptocephali and establishment of a new fish family “Neocyematidae” based on larvae, adults and mitogenomic gene orders. PLoS One. 2018;13: 1–23. doi:10.1371/journal.pone.0199982

66. Rozewicki J, Li S, Amada KM, Standley DM, Katoh K. MAFFT-DASH: Integrated protein sequence and structural alignment. Nucleic Acids Res. 2019;47: W5–W10. doi:10.1093/nar/gkz342

67. Lanfear R, Frandsen PB, Wright AM, Senfeld T, Calcott B. PartitionFinder 2: new methods for selecting partitioned models of evolution for molecular and morphological phylogenetic analyses. Molecular biology and evolution. 2016. doi:dx.doi.org/10.1093/molbev/msw260

68. Lewis PO. A Likelihood Approach to Estimating Phylogeny from Discrete Morphological Character Data. Syst Biol. 2001;50: 913–925. doi:10.1080/106351501753462876

69. Ronquist F, Teslenko M, Van Der Mark P, Ayres DL, Darling A, Höhna S, et al. MrBayes 3.2: Efficient bayesian phylogenetic inference and model choice across a large model space. Syst Biol. 2012;61: 539–542. doi:10.1093/sysbio/sys029

70. Arratia G. Anaethalion and similar teleosts (Actinopterygii, Pisces) from the Late Jurassic (Tithonian) of Southern Germany and their relationships. Paleontographica. 1987.

71. Poyato-Ariza FJ, Wenz S. Naiathaelon okkidion nov.gen. nov. sp., (Teleostei, Elopomorpha) from the Early Tithonian of Canjuers (Var, France). Geobios. 1993; 157–166.

72. Rambaut A, Drummond AJ, Xie D, Baele G, Suchard MA. Posterior summarisation in Bayesian phylogenetics using Tracer 1.7. Syst Biol. 2018. doi:10.1093/sysbio/syy032 Tags:

73. Xie W, Lewis PO, Fan Y, Kuo L, Chen MH. Improving marginal likelihood estimation for bayesian phylogenetic model selection. Syst Biol. 2011;60: 150–160. doi:10.1093/sysbio/syq085

74. Müller J. Ueber den Bau und die Grenzen der Ganoiden, und über das natürliche System der Fische. Archiv für Naturgeschichte. 1845;1: 91–141.

75. Greenwood P, Rosen D, Weitzman S, Myers G. Phyletic studies of teleostean fishes, with a provisional classification of living forms. Bull Am Mus Nat Hist. 1966;131: 339–456. doi:10.1109/23.221050

76. Goodrich E. A Treatise on Zoology. Part IX: Vertebrata Craniata. Lankester R, editor. 1909.

77. Gill T. Three new families of fishes added to the deep-sea fauna in a year. American Naturalist. 1884;18.

78. Linnaei C. Systema Naturæ. 10th ed. Sumptibus Guilielmi Engelmann; 1758.

79. Robins C. The phylogenetic relationships of the anguilliform fishes. In: Bohlke E, editor. Fishes of the Western North Atlantic Volumen One: Orders Anguilliformes and Saccopharyngiformes. Sears Foundation for Marine Research; 1989. pp. 9–24.

80. Broughton RE, Betancur-R. R, Li C, Arratia G, Ortí G. Multi-locus phylogenetic analysis reveals the pattern and tempo of bony fish evolution. PLoS Curr. 2013;5: ecurrents.tol.2ca8041495ffafd0c92756e75247483e. doi:10.1371/CURRENTS.TOL.2CA8041495FFAFD0C92756E75247483E

81. Betancur-R R, Broughton RE, Wiley EO, Carpenter K, López JA, Li C, et al. The Tree of Life and a New Classification of Bony Fishes. PLoS Curr. 2013;5: ecurrents.tol.53ba26640df0ccaee75bb165c8c26288. doi:10.1371/CURRENTS.TOL.53BA26640DF0CCAEE75BB165C8C26288

82. Hughes LC, Ortí G, Huang Y, Sun Y, Baldwin CC, Thompson AW, et al. Comprehensive phylogeny of ray-finned fishes (Actinopterygii) based on transcriptomic and genomic data. 2018;115: 6249–6254. doi:10.5061/dryad

83. Near TJ, Eytan RI, Dornburg A, Kuhn KL, Moore JA, Davis MP, et al. Resolution of ray-finned fish phylogeny and timing of diversification. Proc Natl Acad Sci U S A. 2012;109: 13698–13703. doi:10.1073/PNAS.1206625109

84. Priede IG. Deep-Sea Fishes: Biology, Diversity, Ecology and Fisheries. Cambridge University Press; 2017.

85. Nelson J. Gill arches of teleostean fishes. Ph.D. Thesis, University of Hawaii. 1966.

86. Castle PHJ. Deep-sea eels: Family Synaphobranchidae. Galathea Report. 1964;7: 29–42.

87. Karmovskaya ES, Merrett NR. Taxonomy of the deep-sea eel genus, Histiobranchus (Synaphobranchidae, Anguilliformes), with notes on the ecology of H. bathybius in the eastern North Atlantic. J Fish Biol. 1998;53: 1015–1037. doi:10.1006/jfbi.1998.0765

88. Saldanha L, Merrett NR. The osteology of Ilyophis blachei and its contribution to a diagnosis of the Synaphobranchidae (Pisces, Anguilliformes). Zool J Linn Soc. 1987;91: 343–356. doi:10.1111/j.1096-3642.1987.tb01419.x

89. Svendsen FM, Byrkjedal I. Morphological and molecular variation in Synaphobranchus eels (Anguilliformes: Synaphobranchidae) of the Mid-Atlantic Ridge in relation to species diagnostics. Mar Biodivers. 2013;43: 407–420. doi:10.1007/s12526-013-0168-1

90. Robins C. The Comparative Morphology of the Synaphobranchid Eels of the Straits of Florida. Proc Acad Nat Sci Phila. 1971;123: 153–204.

91. Robins C. Family Synaphobranchidae. In: Bohlke E, editor. Fishes of the Western North Atlantic Part Nine: Orders Anguilliformes and Saccopharyngiformes. Sears Foundation for Marine Research; 1989. pp. 207–253.

92. Johnson J. Descriptions of some new genera and species of fishes obtained at Madeira. Proceedings of the Zoological Society of London 1862 (pt 2). 1862.

93. Aronson R. Foraging behavior of the West Atlantic trumpetfish, Aulostomus maculatus: use of large,herbivorous, reef fishes as camouflage. Bull Mar Sci. 1983;33.

94. Pratchett MS, Hoey AS, Wilson SK, Hobbs J-PA, Allen GR. Habitat-use and Specialisation among Coral Reef Damselishes. In: Fredereich B, Parmentier E, editors. Biology of damselfishes. CRC Press; 2016.

95. Thompson R, Mumot JL. Aspects of the biology and ecology of Caribbean reef fishes: Serranidae (hinds and groupers). J Fish Biol. 1978;12: 115–146.

96. Bellwood DR. The Eocene fishes of Monte Bolca: the earliest coral reef fish assemblage. Coral Reefs. 1996;15: 11–19. doi:10.1007/s003380050025

97. Regan CT. XLIX. — The osteology and classification of the Teleostean fishes of the order Apodes. Annals and Magazine of Natural History. 1912;10: 377–387. doi:10.1080/00222931208693250

98. Regan CT. The Classificatlon of Teleostean Fishes. Annals and Magazine of Natural History. 1909.

99. Vaillant L. Sur un poisson des grandes profondeurs de l’Atlantique, l’Eurypharynx pelecanoides. C R Hebd Seances Acad Sci. 1882;95: 1226– 1228.

100. Inoue JG, Miya M, Miller MJ, Sado T, Hanel R, Hatooka K, et al. Deep-ocean origin of the freshwater eels. Biol Lett. 2010;6: 363–366. doi:10.1098/rsbl.2009.0989

101. Santini F, Harmon LJ, Carnevale G, Alfaro ME. Did genome duplication drive the origin of teleosts? A comparative study of diversification in ray-finned fishes. BMC Evol Biol. 2009;9: 1–15. doi:10.1186/1471-2148-9-194

102. Friedman M. Explosive morphological diversification of spiny-finned teleost fishes in the aftermath of the end-Cretaceous extinction. Proceedings of the Royal Society B: Biological Sciences. 2010;277: 1675. doi:10.1098/RSPB.2009.2177

103. Near TJ, Dornburg A, Eytan RI, Keck BP, Smith WL, Kuhn KL, et al. Phylogeny and tempo of diversification in the superradiation of spiny-rayed fishes. PNAS. 2013;110. doi:10.1073/pnas.1304661110

104. Underwood CJ. Diversification of the Neoselachii (Chondrichthyes) during the Jurassic and Cretaceous. Paleobiology. 2006;32: 215–235. doi:10.1666/04069.1

105. Sorenson L, Santini F, Alfaro ME. The effect of habitat on modern shark diversification. J Evol Biol. 2014;27: 1536–1548. doi:10.1111/jeb.12405

106. Flammensbeck CK, Pollerspöck J, Schedel FDB, Matzke NJ, Straube N. Of teeth and trees: A fossil tip-dating approach to infer divergence times of extinct and extant squaliform sharks. Zool Scr. 2018;47: 539–557. doi:10.1111/zsc.12299

107. Schwarzhans W. Otolithen aus den Gerhartsreiter Schichten (Oberkreide: Maastricht) des Gerhartsreiter Grabens (Oberbayern). Palaeo Ichthyologica. 2010;4. doi:10.13140/2.1.4019.2641

108. Schwarzhans W. Reconstruction of the fossil marine bony fish fauna (Teleostei) from the Eocene to Pleistocene of New Zealand by means of otoliths. With studies of Recent congroid, morid and trachinoid otoliths. Memorie della Società Italiana di Scienze Naturali e del Museo di Storia Naturale di Milano. 2019;46.

109. Schwarzhans W, Lee DE, Gard HJL. Otoliths reveal diverse fish communities in Late Oligocene estuarine to deep-water paleoenvironments in southern Zealandia. New Zealand Journal of Geology and Geophysics. 2017;60: 433–464. doi:10.1080/00288306.2017.1365734

110. Schwarzhans W. The otoliths from the middle Eocene of Osteroden near Bramsche, north-western Germany. Neues Jahrb Geol Palaontol Abh. 2007;244: 299–369. doi:10.1127/0077-7749/2007/0244-0299

111. Bratishko A, Udovichenko M. Fish otoliths from the Early Oligocene of Mangyshlak, Kazakhstan. Neues Jahrb Geol Palaontol Abh. 2013;270: 195–208. doi:10.1127/0077-7749/2013/0366

112. Aguilera O, Schwarzhans W, Moraes-Santos H, Nepomuceno A. Before the flood: Miocene otoliths from eastern Amazon Pirabas Formation reveal a Caribbean-type fish fauna. J South Am Earth Sci. 2014;56: 422–446. doi:10.1016/j.jsames.2014.09.021

113. Schwarzhans WW, Nielsen SN. Fish otoliths from the early Miocene of Chile: a window into the evolution of marine bony fishes in the Southeast Pacific. Swiss J Palaeontol. 2021;140. doi:10.1186/s13358-021-00228-w

114. Schwarzhans W. The otoliths from the Miocene of the North Sea Basin. Weikersheim: Backhuys publishers, Leiden and Margraf Publishers. 2010.

115. Agassiz L. Recherches sur les poissons fossiles. Imprimerie de Petitpierre; 1839.

116. Taverne L, Nolf D. Troisieme note sur les poissons des sables de lede (eocene belge): les fossiles autres que les otolithes. Bull Soc Belge Geol; Bel; Da 1978 Publ 1979; Vol 87; No 3-4; P:125-152; Abs Eng; Bibl 2 P; 26 Ill;La. 1978; 125-152.

117. Young SVT, Williams RJ. New information on the cranial anatomy of the eel genus Echelus Rafinesque, 1810 (Ophichthidae: Anguilliformes) from the Early Eocene. Geol Soc Spec Publ. 2008;295: 311–336. doi:10.1144/SP295.15

118. Girard C. Notes upon various new genera and new species of fishes, in the museum of the Smithsonian Institution, and collected in connection with the United States and Mexican boundary survey: Major William Emory, Commissioner. Proceedings of the Academy of Natural Sciences of Philadelphia. 1858;10: 167–171.

119. Gray J. Description of twelve new genera of fish, discovered by Gen. Hardwicke, in India, the greater part in the British Museum. Zoological Miscellany. 1831; 7–9.

120. Schwarzhans W. Otoliths from casts from the Eocene Lillebælt Clay Formation of Trelde Næs near Fredericia (Denmark), with remarks on the diet of stomatopods. Neues Jahrb Geol Palaontol Abh. 2007;246: 69–81. doi:10.1127/0077-7749/2007/0246-0069

